# Microtubule plus-end dynamics link wound repair to the innate immune response

**DOI:** 10.1101/512632

**Authors:** Clara Taffoni, Shizue Omi, Caroline Huber, Sebastien Mailfert, Matthieu Fallet, Jonathan J. Ewbank, Nathalie Pujol

## Abstract

As a first line of defence against the environment, the epidermis protect animals from infection and physical damage. In *C. elegans*, wounding the epidermal epithelium triggers both an immune reaction and a repair response. Exactly how these are controlled, and the degree to which they are inter-connected remains unclear. To address these questions, we established a simple system for simultaneously inflicting precise laser wounds and imaging at high spatial and temporal resolution. We show that in *C. elegans*, wounding provokes a rapid sealing of the plasma membrane, involving reorganisation of phosphatidylinositol 4,5- bisphosphate domains. This is followed by a radial recruitment at the wound site of EBP-2/EB1, a protein that binds the plus ends of microtubules. EB1 recruitment is accompanied by a reorganisation of microtubules, required for the subsequent recruitment of actin and wound closure. It is also required for the directed trafficking towards the site of injury of the key signalling protein SNF-12. In the absence of SNF-12 recruitment, there is an abrogation of the immune response. Our results suggest that microtubule dynamics coordinate the cytoskeletal changes required for wound repair and the concomitant activation of the innate immune response.

## Introduction

Inducible immune responses are ubiquitous features of animal defences against infection. In one well-studied example, when the fungus *Drechmeria coniospora* infects *Caenorhabditis elegans*, by piercing the cuticle and growing through the underlying epidermis, it triggers an innate immune response, characterized by the induction of expression of a battery of antimicrobial peptide (AMP) genes (Pujol et al., 2008b; Taffoni and Pujol, 2015). These include the genes of the *nlp-29* cluster (Pujol et al., 2012), which are principally controlled in a cell autonomous manner (Lee et al., 2018). Sterile wounding also upregulates the expression of the *nlp-29* cluster (Pujol et al., 2008a). In both cases, these changes in gene expression result from the activation of the GPCR DCAR-1 that acts upstream of a conserved p38 MAPK signalling cassette (Zugasti et al., 2014). The p38 MAPK PMK-1 itself acts upstream of the STAT transcription factor-like protein STA-2. Although the precise details are as yet unclear, activation of STA-2 is believed to require the SLC6 family protein SNF-12 (Dierking et al., 2011).

Triggering the immune response upon wounding has a prophylactic role, and is a safeguard against the potential entry of pathogenic microbes. To ensure longer term protection, a wounded tissue must also be repaired to maintain organismal integrity. Thus in addition to AMP gene induction, skin wounding in *C. elegans* also triggers a Ca^2+^-dependent signalling cascade that promotes wound repair through actin ring closure (Xu and Chisholm, 2011). The nematode skin is mainly a syncytium, except at the head and tail, a single multinucleate epidermal cell, hyp7, covers the whole animal (Altun and Hall, 2014). Its repair processes exhibit some of the characteristics previously described for single cell wound repair in other organisms (Sonnemann and Bement, 2011). On the other hand, actin ring closure in *C. elegans* does not rely on myosin-II contractility and the “purse string” mechanism described in other organisms, but instead requires Arp2/3-dependent actin polymerization.

A small number of reports have indicated that MTs cooperate with the actomyosin cytoskeleton during wound closure. In a single-cell wound-healing model using *Xenopus laevis* oocytes, MTs were shown to be required to restrict the assembly zone of actin and myosin around the wound edge (Bement et al., 1999; Mandato and Bement, 2003). A study in which *D. melanogaster* embryos were used also revealed that perturbation of MT dynamics, in a + end-binding protein 1 (EB1) mutant, resulted in a delay of actomyosin assembly at the wound edge in multi-cellular wounds (Abreu-Blanco et al., 2012). The direct role of MTs in the *C. elegans* wound response has not been previously addressed.

The tissue repair processes in *C. elegans* have been described to act in parallel to the innate immune response that accompanies wounding (Xu and Chisholm, 2011). Several pieces of evidence suggest, however, a coordination of the two responses. Earlier work had shown that loss of DAPK-1, homolog of death-associated protein kinase, causes both undue tissue repair and an inappropriate activation of an immune response (Tong et al., 2009). A more recent study showed that loss of *dapk-1* function causes excessive microtubule (MT) stabilization (Chuang et al., 2016). These results suggest that MT stability could directly influence immune signalling.

Here, we develop the use of a 405 nm laser to wound the *C. elegans* epidermis and characterise the subsequent subcellular events in vivo. We show for the first time a rapid membrane reorganisation of phosphatidylinositol 4,5-bisphosphate (PIP_2_) domains, as well as a recruitment of EB1 and reorganisation of MTs. We demonstrate that specific inactivation in the adult epidermis of several alpha and beta tubulin isotypes leads to the abrogation of the immune response upon wounding or infection. Indeed, tubulin inactivation leads not only to decreased recruitment of actin at the wound site, but also of the key signalling protein SNF-12 and a block of the subsequent immune response. These results suggest that recruitment and stabilisation of MTs to wounds coordinate the activation of the innate immune response and wound repair.

## Results

### The *C. elegans* epidermis can be wounded with a 405 nm laser

The adult epidermis in *C. elegans* is a single cell layer, covered on its apical surface by an impermeable cuticle that serves as an exoskeleton. The epidermal syncytium hyp7 can be divided into two distinct regions, one lateral, where the nuclei and most of the cytoplasm are, and the dorso-ventral part above the muscles where the apical and basolateral plasma membranes of hyp7 are juxtaposed and crossed by hemidesmosomes that anchor the muscles to the cuticle exoskeleton (Fig. 1A) (Altun and Hall, 2014). The large size of hyp7 make it an attractive model for the study of cellular wound repair mechanisms (Pujol et al., 2008a; Xu and Chisholm, 2011). Previous studies on *Xenopus* oocytes have demonstrated that the plasma membrane can be perforated in an extremely targeted manner using standard 405 nm microscopy imaging lasers (Burkel et al., 2012; Mandato and Bement, 2001). We found that we could efficiently wound the hyp7 lateral syncytial epidermis of *C. elegans* (Fig. 1A) with a short pulse (1-3 s) of 405 nm laser light in Fluorescence Recovery After Photobleaching (FRAP) mode (22 mW). Coupled with spinning disc microscopy, this provides a reproducible way to monitor *in vivo* the immediate consequences of a carefully controlled injury and the subsequent steps in the wound healing process. Using this system, we first confirmed all the previously described wound hallmarks (Pujol et al., 2008a; Xu and Chisholm, 2011), such as an immediate autofluorescent scar (Fig. S1A), a Ca^2+^ burst (Fig. 1B), the formation of an actin ring (Fig. S1B) and later the induction of AMP reporter gene expression (Fig. S1C).

**Fig. 1:**
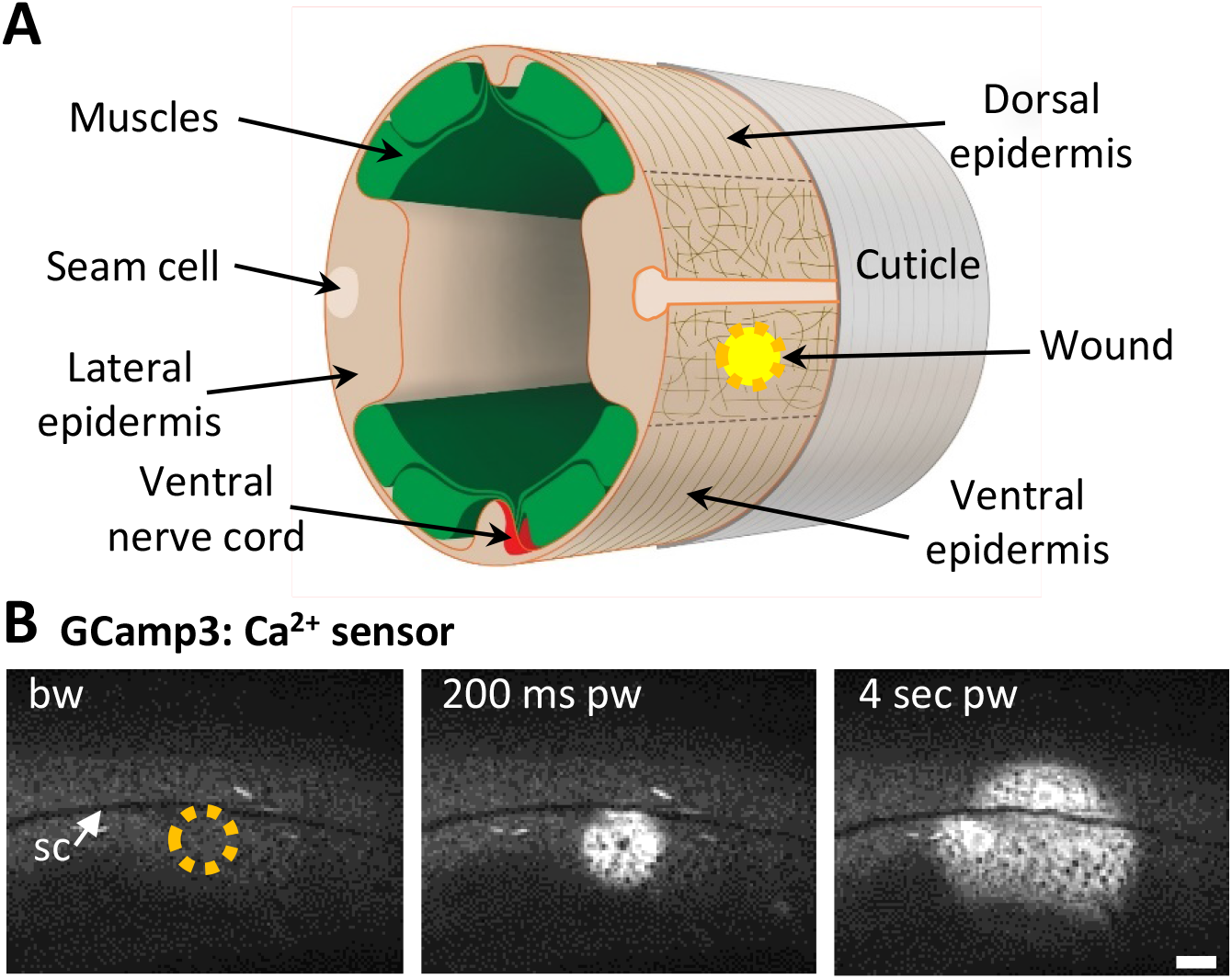
Using a 405 nm laser to wound the worm epidermis. (A) Schematic view of a section through an adult *C. elegans* worm near the mid-body. The main syncytial epidermis, hyp7 (buff), with its adjacent cells (seam cells in pink, nerve cord in red and muscles in green) can be divided into 2 main regions, the lateral epidermis (delimited by the dashed black line) and the thin dorso-ventral epidermis where muscles are anchored to the cuticle. Microtubules (thin taupe lines) are disordered on the apical surface of the lateral epidermis. The typical position of a wound is indicated by the yellow circle. Figure adapted from one kindly provided by Christopher Crocker, WormAtlas (Altun and Hall, 2014). (B) Wounding the lateral epidermis with a 405 nm laser causes a rapid increase in intracellular Ca^2+^, measured using GCamp3. Representative green fluorescence images from a worm carrying a *col-19p::GCamp3* reporter transgene. Here and in subsequent images, the dashed circle is centred on the wound site; bw, before wound (200 ms); pw, post-wound; sc, seam cells. Scale bar 10 μm.

### Membrane is rapidly reorganised at the wound site

In *Xenopus* oocytes, a wound is rapidly patched through mobilisation and recruitment of membrane from a local pool of vesicles. We could similarly observe rapid membrane recruitment at the wound site in the epidermal syncytium using a strain expressing a prenylated form of GFP (*dpy-7p::GFP::CAAX*). Within 15 sec after wounding, large heterogeneous membrane domains regrouped and extended towards the wound (Fig. 2A, Movie 1). PIP_2_ has been found segregated into distinct membrane pools in the plasma membrane in other species, consistent with its wide range of cellular functions including regulating the adhesion between the actin-based cortical cytoskeleton and the plasma membrane (Raucher et al., 2000). We found that the plasma membrane of the hyp7 syncytium was also heterogeneously labelled using a strain in which PIP_2_ was specifically labelled (*wrt-2p::GFP::PH-PLC1δ* (Wildwater et al., 2011)). Upon wounding, we observed a rapid reorganization of these membrane domains around the wound (Fig. 2B, Movie 2). These processes are likely to provide an immediate barrier to potential leakage of cellular components as previously proposed in other systems (Davenport et al., 2016; Vaughan et al., 2014).

**Fig. 2:**
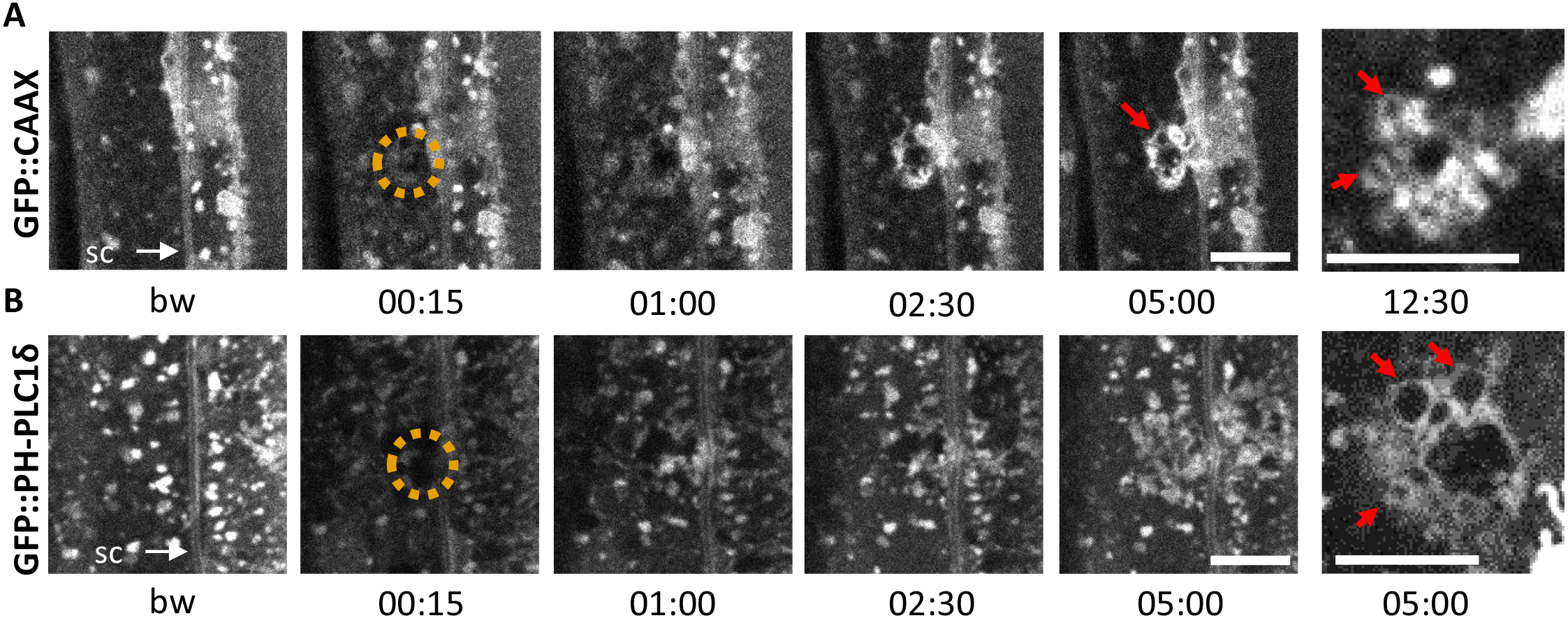
Membrane and lipid domain reorganization at the wound site. Upon laser wounding, membrane and lipids are rapidly recruited to the wound site. The red arrows highlight examples of vesicle fusion. Representative images of worms carrying *dpy-7p::GFP::CAAX* (A) and *wrt-2p::GFP::PH-PLClδ* transgenes (B). bw, before wound; time post-wound [min:sec]; sc, seam cells. Scale bar 5 μm.

### Actin and MTs reorganize at the wound site forming concentric rings

Wounding the *C. elegans* epidermis causes cytoskeleton rearrangements, with an actin ring forming at the wound site (Xu and Chisholm, 2011). We wanted to monitor the dynamics of actin recruitment and so constructed a strain in which the filamentous actin binding peptide Lifeact (Riedl et al., 2008), labelled with mKate, was specifically expressed in the adult epidermis (*col-62p::Lifeact::mKate*) (Fig. 3A). Under resting condition in the young adult, in the lateral epidermis, we observed a disorganised dot-like pattern, with only occasional fine actin filaments (Fig. 3A, bw (before wound)). This is in contrast to the highly organised circumferential bundles of actin found in the same hyp7 cell above the muscles, as previously described (Costa et al., 1997; Lazetic et al., 2018). Upon wounding, while the pre-existing actin filaments appeared stable, actin was recruited to the wound site within 2-4 minutes and formed a well-defined actin ring after 5 minutes. This subsequently constricted as the wound closed (Fig. 3A and Movie 3), in line with previous observations (Chuang et al., 2016).

**Fig. 3:**
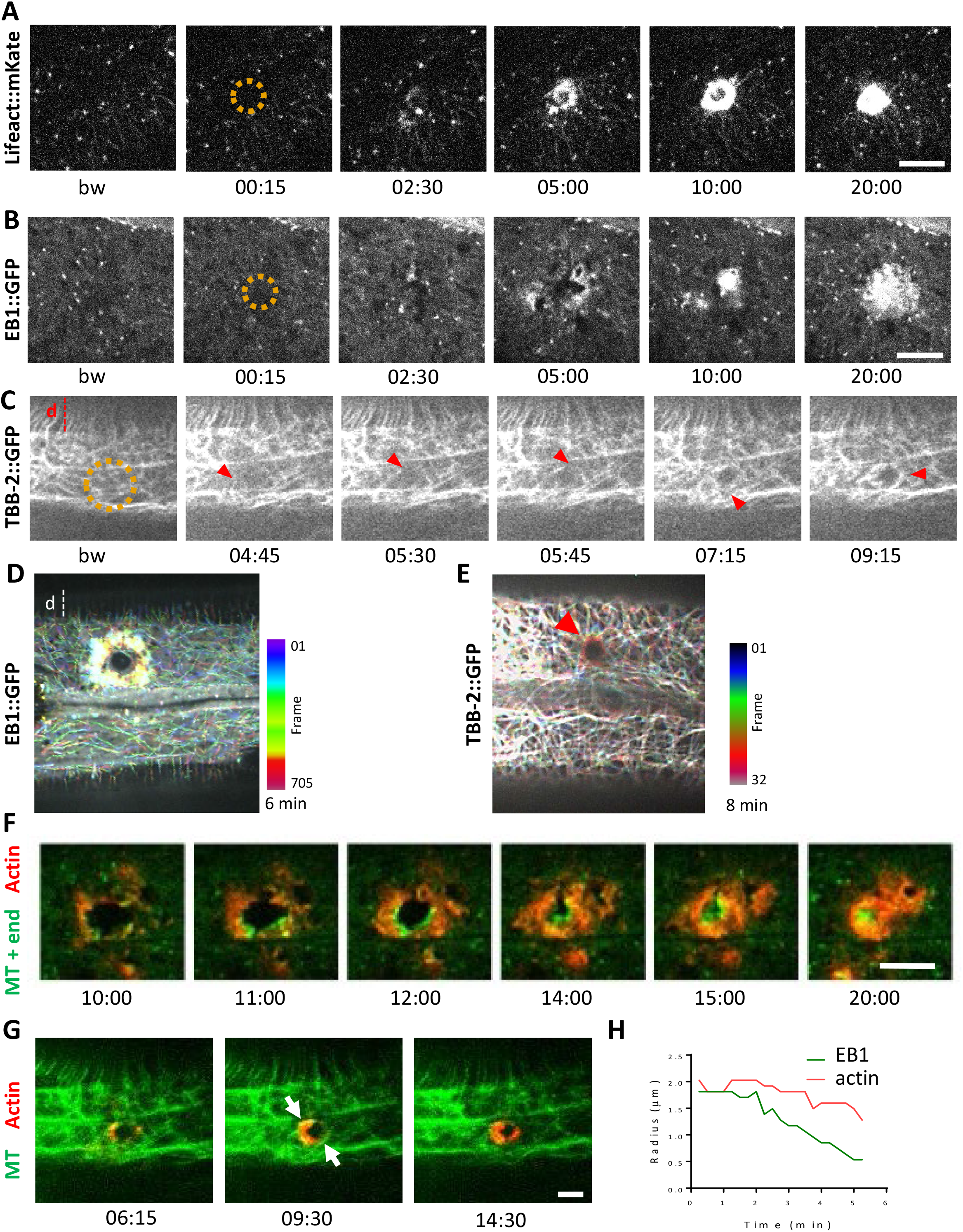
Cytoskeleton reorganization at the wound site in the lateral epidermis. (A) In unwounded worms, in the lateral epidermis actin is sparsely structured. Laser wounding causes actin recruitment (as early as 2.5 min) and actin ring formation (clearly visible by 5 min). These rings close with time. Representative images of a strain carrying a *col-62p::Lifeact::mKate* reporter. Laser wounding also causes EB1 recruitment and MT reorganization (highlighted with red arrows) at the wound site as visualized in strains carrying *col-19p::EBP-2::GFP* (B) and *TBB-2::GFP* (C), respectively; d, dorsal epidermis. After wounding, EB1 (D) and MT (E, highlighted with red arrow) reorganise around the wound as represented with a colour coded temporal projection (0.2 s or 15 s frame interval, respectively). (F-G) MT congregation precedes actin accumulation. Simultaneous visualization of MT (green) and actin (red) using strains carrying EB1::GFP (F) or TBB-2::GFP (G) together with *col-62p::Lifeact::mKate*. The white arrows indicate representative MT bundles. bw, before wound; time post-wound [min:sec]; scale bar 5 μm. (H) EB1 (green) ring contraction precedes that of actin (red) during wound closure. Radial profile where the radius between the maximum intensity is represented over time for EB1 in green and actin in red.

Microtubules (MTs) are another essential component of the cytoskeleton, but their role in wound healing has not previously been addressed directly in *C. elegans*. To look at MT dynamics, we imaged worms expressing a tagged form of EB1/EBP-2, a protein that binds to the + end of growing MTs (*col-19p::EBP-2::GFP*; EB1::GFP for brevity). Under resting conditions EB1::GFP exhibited rapid, comet-like movement throughout hyp7, with a speed (0.26 ± 0.10 μm/sec, Table 1) close to that determined in a recent study (0.3 μm/sec) (Chuang et al., 2016). Upon wounding, EB1::GFP comets accumulated within 2 min around the wound site (Fig. 3B and Movie 4). This was concomitant with a reduction of their average speed in the vicinity of the wound, to 0.16 ± 0.06 μm/sec (Table 1), indicative of a local stabilisation of MTs.

**Table 1.** Reporter protein dynamics in the epidermis

| Type of Dynamics | Transgene | Velocity (mean ± SEM) $\mu\text{m s}^{-1}$ | |
| --- | --- | --- | --- |
|  |  | before wound | after wound |
| MT plus-end growth | *col-19p::EBP-2::GFP* | 0.26 ± 0.10 $\mu\text{m/sec}$<br>(n=33) | 0.16 ± 0.06 $\mu\text{m/sec}$<br>(n=33) |
| endosome transport | *dpy-7p::GFP::RAB-11* | 1.31 ± 0.33 $\mu\text{m/sec}$<br>(n=50) | 1.14 ± 0.47 $\mu\text{m/sec}$<br>(n=40) |
| SNF-12 | *col-12p::SNF-12::GFP* | immobile* | 0.007 ± 0.004 $\mu\text{m/sec}$<br>(n=40) |

\*less than 1% of SNF-12 clusters move with a speed of 0.017 ± 0.005 $\mu\text{m/sec}$ (n=21)

To confirm that EB1 dynamics reflected local MT reorganization at the wound site, we used 2 independent strains where one of the 16 *C. elegans* tubulin proteins, TBB-2, was labelled with GFP (*Si [col-19p::GFP::TBB-2]* and *KI [tbb-2p::TBB-2::GFP]*; here termed TBB-2::GFP). We observed that MTs were arranged essentially in a longitudinal but haphazard manner in the lateral epidermis, in marked contrast to their parallel and circumferential organisation in the dorso-ventral epidermis, over the muscle quadrants (Fig. 3C), as previously described (Costa et al., 1997; Wang et al., 2015). The MTs in the lateral epidermis were highly dynamic. Consistent with our observations using EB1::GFP, after wounding they locally regrew to surround the site of injury (Fig. 3C and Movie 5). The dynamic recruitment at the wound site of EB1::GFP and TBB-2::GFP was visualised with a temporal projection (Fig. 3D-E), confirming that EB1 massive recruitment is linked to growth of MT around the wound. To visualise simultaneously actin and MTs dynamics, we generated two strains, each containing two reporter transgenes, either EB1::GFP or TBB-2::GFP with Lifeact::mKate. We observed that actin and MTs formed partially overlapping concentric rings and that the EB1::GFP ring was circumscribed by actin (Fig. 3F-G and Movie 6 & 7). As assayed with EB1::GFP, MTs grew in front of the actin ring as it closed (Fig. 3H). These results are consistent with MT reorganisation driving actin ring formation.

### *tba-2* or *tbb-2* inactivation results in abrogation of EB1 dynamics in the epidermis

To test whether MT growth at the wound site does promote actin polymerization, we sought a way to block MT dynamics prior wounding the epidermis. Due presumably to their impermeable cuticle, treatment of adult worms with commonly used drugs, such as colchicine, was either ineffective or had irreproducible effects in our hands. Although inactivating various genes that affect cuticle formation can increase permeability and so render *C. elegans* more susceptible to drugs (Loer et al., 2015; Partridge et al., 2008), this uniformly affects the epidermal innate immune response (Dodd et al., 2018; Zugasti et al., 2016). We therefore used RNAi to interfere directly with tubulin gene expression. Since eliminating MTs provokes developmental delay and lethality, worms were subjected to a relatively brief (24 h) RNAi treatment from the L4 stage. Under these conditions, worms completed their development normally and exhibited no overt morphological defects. We chose one tubulin alpha *tba-2* and one tubulin beta *tbb-2*, as these genes have been reported to be highly expressed in the epidermis (Cao et al., 2017; Hutter and Suh, 2016). In young adult *tba-2*(RNAi) or *tbb-2*(RNAi) worms, EB1::GFP comets were no longer visible suggesting that MT dynamics had been severely compromised (Fig. 4A and 4B and Fig. S2A). Interestingly, this effect was specific to the epidermis (hyp7) since highly motile comets were still visible in the seam cells (Fig. 4B). Upon wounding, EB1::GFP was not recruited to the wound site (Fig. 4A and 4C). This suggests that constant expression of *tba-2* and *tbb-2* genes is required for the normal dynamic behaviour of the non-centrosomal MTs in the epidermal syncytium as well as the redistribution of MTs caused by injury.

**Fig. 4:**
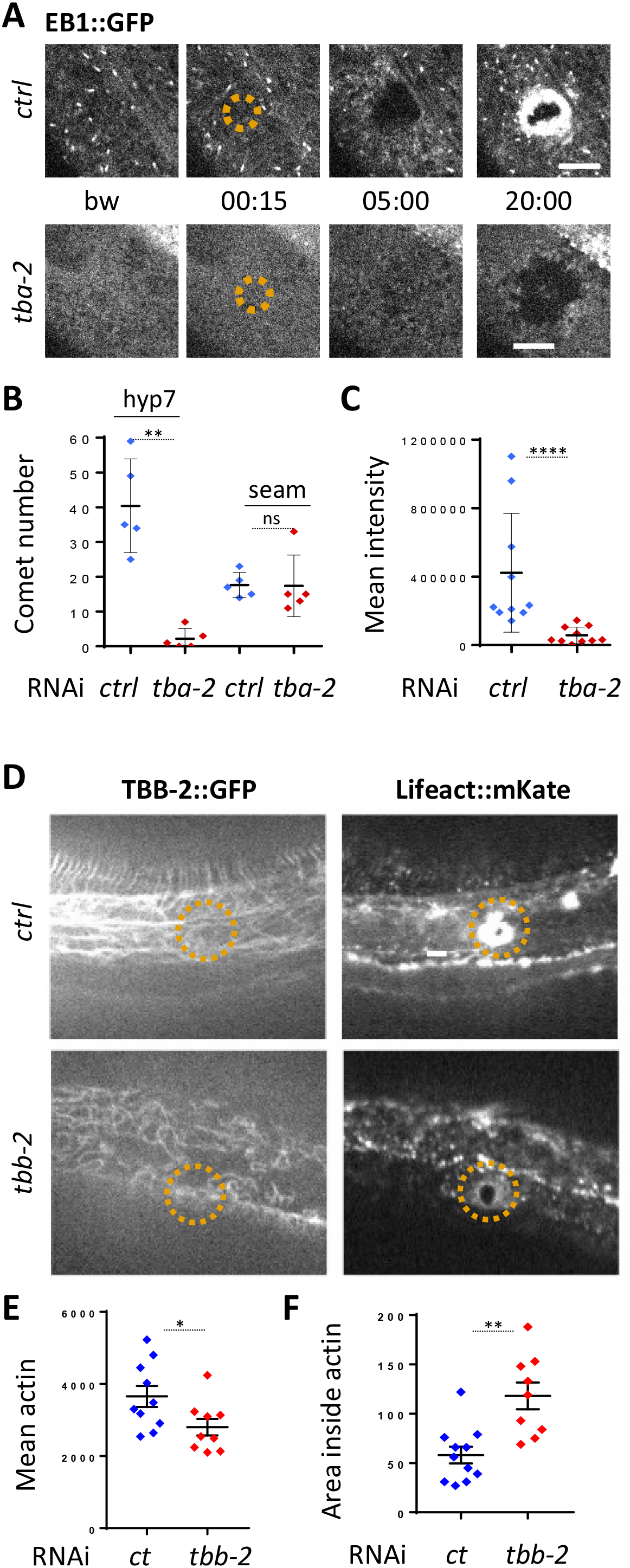
TBA-2 is a tubulin required for non-centrosomal microtubule dynamics in the lateral epidermis. (A) TBA-2 is required for EB1 comet dynamics and EB1 recruitment at the wound site. Representative images of *col-19p::EBP-2::GFP* in control (*ctrl*) and *tba-2* RNAi treated worms; time post wound [min:sec]; scale bar 5 μm. (B) Quantification of EB1 comet number in either lateral epidermis or seam cells (n = 5 worms) before wounding. (C) Quantification of EB1 recruitment at the wound site, at 10 min post wounding (n = 10 wounds). ns p > 0,05, ** p < 0.01 and **** p < 0.0001; non parametric Mann-Whitney test. Representative images of TBB-2::GFP (left) and Lifeact::mKate (right) (D) and quantification of the intensity of actin (E) and the area inside the actin ring (F) in control and *tbb-2* RNAi treated worms 15 mn after wounding; * p < 0.05, ** p < 0.01 non parametric Mann-Whitney test.

### Abrogation of MT dynamics decreases actin recruitment at the wound site

To address the question whether MTs are required for the formation of the actin ring, we analysed actin ring formation upon wounding in *tbb-2*(RNAi) worms. In resting conditions in young adult worms, short abrogation of *tbb-2* altered the MT network; it became less dense and less longitudinally structured in the lateral epidermis (Fig. 4D). Upon wounding these worms, less actin was recruited around the wound (Fig. 4D-E). Moreover, the actin ring did not constrict as quickly as in wild type worms, as measured by the area inside the ring after 15 min (Fig. 4D & F). Interestingly, the relationship between MTs and actin was not reciprocal. Thus reduction of the level of actin via inactivation of *act-5* did not change the overall pattern of MTs nor EB1 dynamics (FigS2 A&B). This suggests that the dynamic reorganisation of MTs around the wound is indeed important for actin ring closure and might helps driving it.

### Microtubule dynamics is required for the immune response

We had previously demonstrated in a genome-wide RNAi screen that knocking down certain genes associated with MT function, including *tba-2* and *tba-4*, leads to an abrogation of the induction of the AMP gene reporter *nlp-29p::GFP* usually caused by *Drechmeria coniospora* infection (Table S1, (Zugasti et al., 2016)). We found more precisely that targeted and short inactivation just before the adult stage of these 2 tubulin α genes led to a block of AMP induction after infection in the adult (Fig. 5A). In addition, we showed that *tba-2* and *tba-4* are also required for AMP reporter gene induction after wounding (Fig. 5A). MTs are formed by dimers of tubulin, each consisting of two subunits, tubulin α and tubulin β. In the *C. elegans* genome, there are 9 α (TBA) and 6 β (TBB) genes. Some have been shown to act in specific cells or tissue, such as *mec-7* (β) and *mec-12* (α) in the mechanosensory cells (Savage et al., 1989), other are predicted to be ubiquitously expressed, and several to be expressed in the epidermis (Harris et al., 2010). We extended our investigation to all tubulin genes by assaying the effect of individually knocking down their expression by RNAi specifically in the adult epidermis. In addition to *tba-2* and *tba-4*, we found that inactivation of tubulin β genes *tbb-1* and *tbb-2* was also associated with a block of AMP reporter gene induction upon infection (Fig. 5B-C). We found that this was not a consequence of reduced spore binding to the worm cuticle (Fig. S3). Using the TBB-2::GFP strain, we also showed that inactivation of the genes that blocked the immune response altered the pattern of MTs, confirming the expression and functional importance of these isoforms in the adult epidermis (Fig. S2C).

**Fig. 5:**
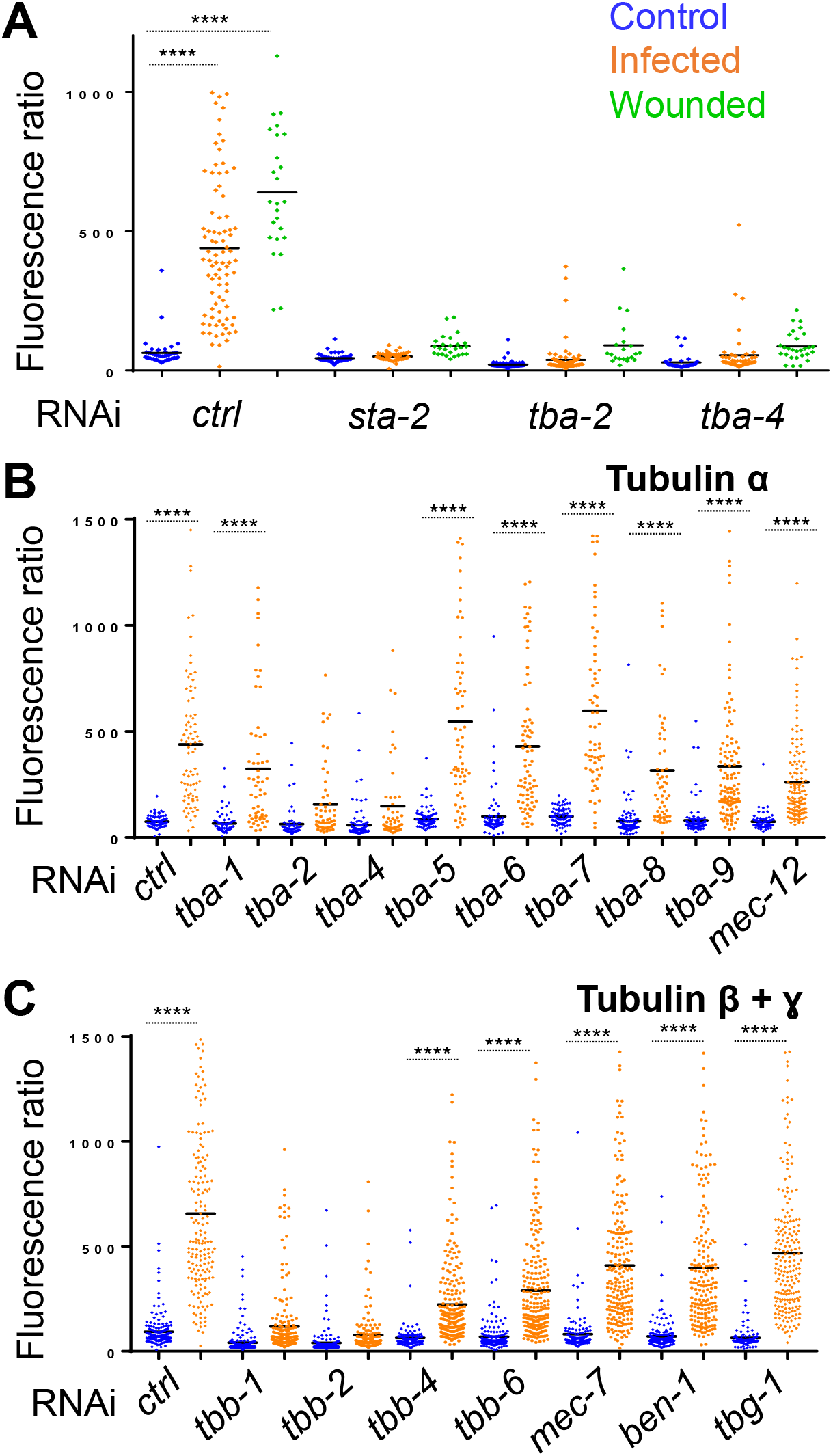
Specific tubulin isoforms are involved in the activation of the immune response upon wounding and fungal infection. Quantification of relative green fluorescence in a strain carrying the *nlp-29p::gfp* transcriptional reporter after RNAi against different tubulin α, β et γ genes in non-infected worms or after infection with *D. coniospora* or after wounding (blue, orange and green symbols, respectively). (A) Worms were fed on RNAi bacteria from the L4 stage and after 24 h infected or wounded; sta-2(RNAi) is known to block the immune response (Dierking et al., 2011). (B, C) Worms sensitive to RNAi principally in the adult epidermis were fed on RNAi clones from L1 stage and infected with *D. coniospora* at the young adult stage. Mean are represented in black, number of worms in 5A: 47, 87, 25, 48, 46, 26, 66, 107, 21, 33, 52, 28; in 5B: 106, 82, 67, 59, 63, 57, 112, 51, 95, 65, 88, 76, 79, 65, 128, 58, 98, 121, 69, 138; in 5C: 228, 180, 199, 210, 201, 162, 145, 232, 220, 224, 147, 207, 196, 188, 133, 209. Only **** p < 0.0001 is presented; ANOVA Bonferoni’s test. Graphs are representative of the results obtained from at least three independent replicates.

On the other hand, a targeted short inactivation of tubulin-γ (TBG-1) expression did not affect AMP reporter gene induction nor the overall MTs pattern (Fig. 5C and Fig S2C). Since tubulin-γ nucleates MTs at their minus end (Quintin et al., 2016; Wang et al., 2015), this suggests that unlike MT + ends, −ends are less dynamic in the adult epidermis and are not required for the immune response upon wounding. Thus, independent of their role in the maintenance of epidermal structure, and in addition to a function in mediating wound closure upon wounding, MT + end dynamics appear to be required for the regulation of the transcriptional response to injury and fungal infection in the epidermis.

### SNF-12, RAB-5 and RAB-11 get locally recruited upon wounding

We hypothesised that microtubules could be required for controlling the subcellular localization of key signalling molecules that are necessary for the induction of the epidermal immune response. The SLC6 family member SNF-12 was a prime candidate, as we have previously shown that it is localized in clusters in the apical surface of the hyp7 syncytium (Dierking et al., 2011). We therefore characterised further the localization of SNF-12 and its behaviour following wounding. We performed colocalization analysis between SNF-12::GFP and available vesicular and plasma membrane markers (early, late and recycling endosomes, lysosomes, plasma membrane lipids). We did not, however, detect any colocalization (Fig. 6A-B, Fig. S4). On the other hand, we observed that SNF-12 clusters were in the same focal plane as early, late and recycling endosomes (Fig. 6A, Fig. S4 & S5) and PIP_2_ (Fig. 6B). This indicates that SNF-12 is in a yet-to-be defined apical membrane compartment. Interestingly, in the dorso and ventral epidermis, SNF-12 was found in a banded pattern, like the highly organised circumferential cytoskeleton (Fig. 6C, Movie 7). Compared to EB1 and RAB-11, see below, the majority of SNF-12 clusters were static or just vibrating, with only very few moving longer straight distances, presumably along MT tracks, with a low speed of 0.017 ± 0.005 μm/sec (Fig. 6D, Movie 8, Table 1).

**Fig. 6:**
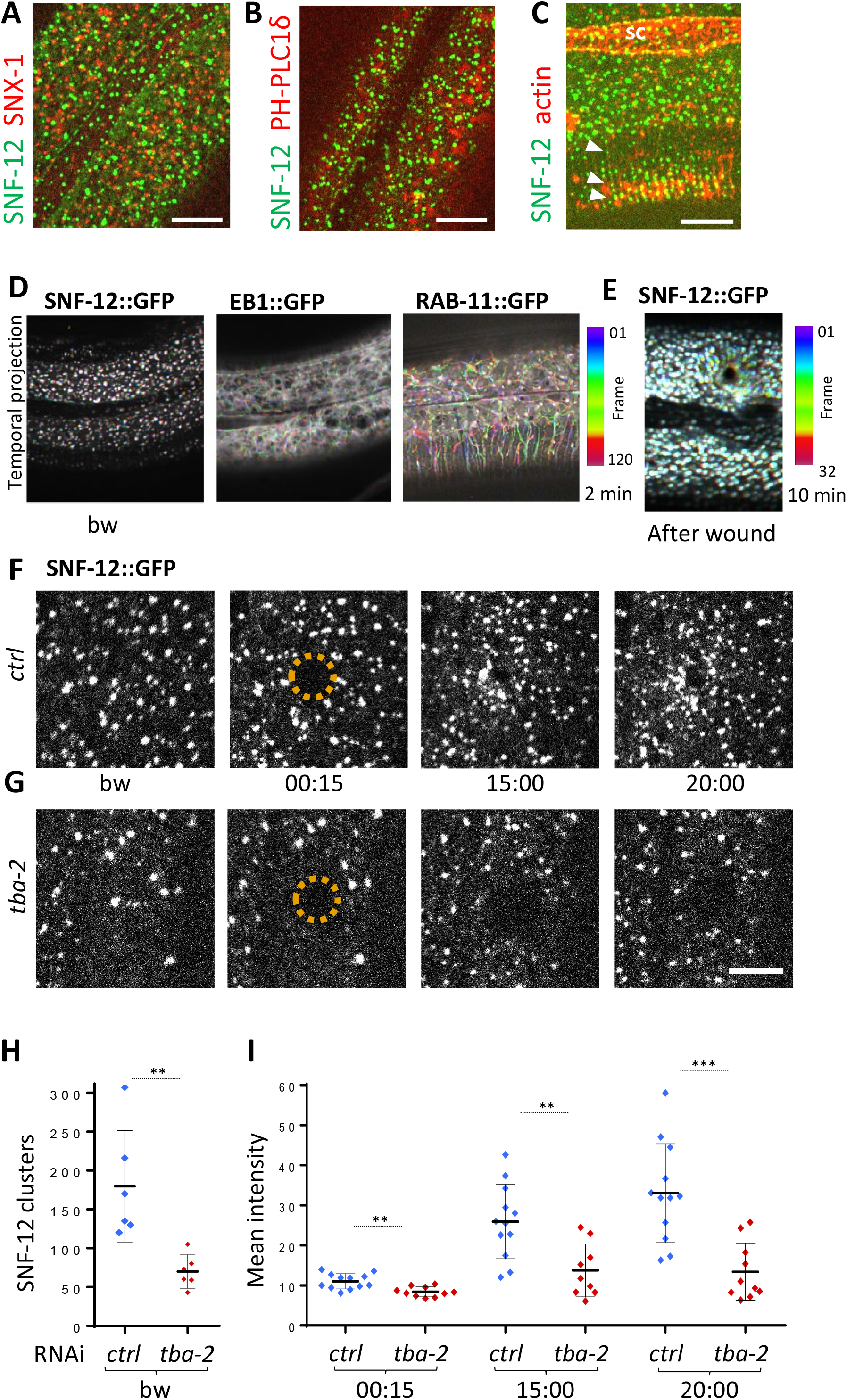
SNF-12 localizes to apical clusters that are recruited at the wound site in a MT-dependent way. Representative images of worms carrying *col-12p::SNF-12::GFP* as well as a red marker of early endosome (*snx-1p::mRFP::SNX-1*; A), membrane lipids (PIP_2_; *ced-1p::mCherry::PH-PLC1δ;* B) or actin (*col-62p::Lifeact::mKate*; C); scale bar 10 μm. (D) The different dynamics of SNF-12, EB1 and RAB-11 are represented with a temporal color coded projection of 120 frames over 2 mn (1 fps) before wounding. (E) After wounding, SNF-12::GFP clusters move toward the wound as represented with a color coded temporal projection of 32 frames over 10 min before wounding and 40 after, 6.30 min after wounding. (F-G) The SNF-12 recruitment to the wound site seen in control animals (F) is abrogated upon *tba-2* RNAi (G); time post wound [min:sec]; scale bar 5 μm. (H) Quantification of SNF-12 cluster number in the lateral epidermis (worm n= 6). (I) Quantification of SNF-12 recruitment at the wound site at different time points post wounding (wound n= 12 for control worms and 10 for *tba-2* RNAi). ** p < 0.01 and *** p < 0.001; non parametric Mann-Whitney test.

When we wounded worms carrying the SNF-12::GFP reporter, we observed a progressive recruitment of SNF-12 to the wound site (Fig. 6E-F, Movie 9), with the majority of the clusters around the wound moving in a directed manner at a low speed of 0.007 ± 0.004 μm/s (Table 1). A similar recruitment was also seen in several other strains containing different tagged forms of SNF-12 (Fig. S6) suggesting that it reflects the behaviour of the endogenous protein. When we disrupted MTs, via RNAi of *tba-2*, SNF-12 patterning and recruitment were severely compromised (Fig. 6G-I). Similarly, both MT and SNF-12 patterning and dynamics upon wounding were also affected when the MT severing protein SPAS-1 was overexpressed in the worm epidermis (Fig. S7). Together, these results suggest that MTs play an important role in SNF-12 localization and dynamics and thereby in the induction of AMP gene expression.

The behaviour of SNF-12 was unusual as in almost all cases when we wounded strains in which vesicles markers were fluorescently tagged, there was no such recruitment (Fig. S5). Thus the movement of SNF-12 towards the wound site was not simply a reflection of cytoskeleton rearrangements or cytoplasmic flow. Two exceptions were RAB-5, a marker of early endosomes, and RAB-11. RAB-5::GFP appeared localised in large static donut-shaped structures (0.3-0.8 μm) arranged in a reticular pattern, exchanging with smaller vesicles (0.2-0.4 μm), as described previously (Chuang et al., 2016) (Fig. 7A, Movie 10). Upon wounding these small vesicles accumulated around the wound site within 10-15 min (Fig. 7A, Movie 11). RAB-11::GFP was found in extremely motile (1.31 ± 0.33 μm/s) vesicles (0.2-0.4 μm) (Table 1). They moved long distances in one direction, presumably on microtubule tracks (Fig. 6D, Movie 12). Upon wounding they started to accumulate around the wound site within 10-15 min (Fig.7B, Movie 13) and reached a maximum density between 30 and 60 min post wounding. Notably, we previously demonstrated that both *rab-5* and *rab-11* are required non-redundantly for the epidermal innate immune response to infection (Dierking et al., 2011; Zugasti et al., 2016). Together, these results suggest that the innate immune response requires coordinated recruitment of both endosomal vesicles and the uncharacterised SNF-12-associated clusters, which is in turn dependent upon MT reorganisation.

**Fig. 7:**
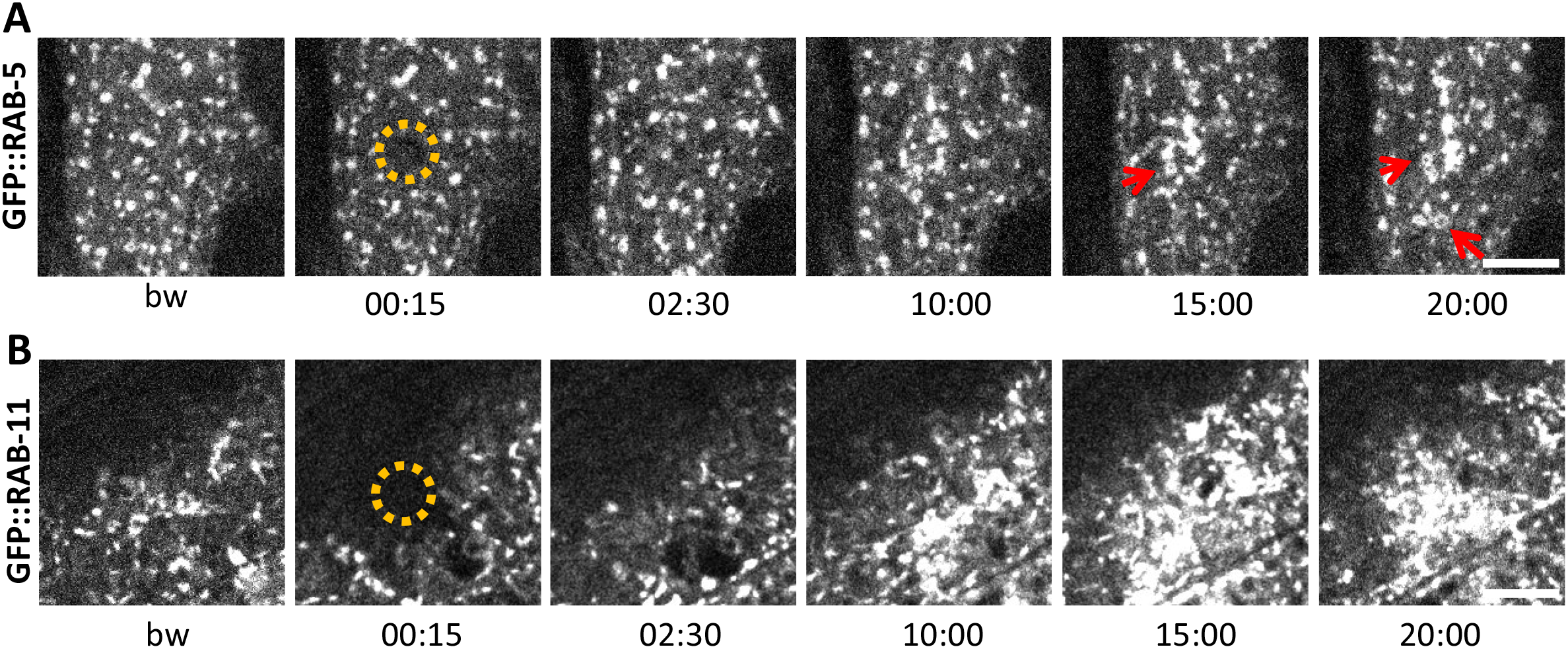
Early and recycling endosomes are recruited to the wound site. Representative images of worms carrying *dpy-7p::GFP::RAB-5* (A) and *dpy-7p::GFP::RAB-11* (B). A fraction of the early endosome marker RAB-5 localizes to donut-shaped structures (red arrows). Upon laser wounding these RAB-5-positive structures (A) as well as RAB-11-positive recycling endosomes (B) are recruited to the wound site within 10 min; time post wound [min:sec]; scale bar 5 μm.

## Discussion

Previous studies of wound healing in *C. elegans* used either imprecise and irreproducible mechanical injury, with a needle, or sophisticated laser rigs not accessible to many laboratories. We demonstrated that a simple FRAP laser can be used to induce wounds with high precision, and can be combined easily with fast image acquisition. Using this system, we observed in the *C. elegans* epidermis the phenomenon of rapid membrane reorganisation previously described in *Xenopus* (Davenport et al., 2016) suggesting that these initial steps of plasma membrane repair, as well as subsequent steps in wound healing (Nakamura et al., 2018), may be conserved throughout Metazoans.

In addition to the acute patching of the membrane, wounding was followed by reorganisation of PIP_2_ domains. PIP_2_ is known to influence the adhesion between the actin-based cortical cytoskeleton and the plasma membrane (Raucher et al., 2000). Thus, its redistribution could be linked to the dramatic changes in the actin cytoskeleton that occurred in the minutes after wounding. The *C. elegans* epidermis is bounded on its apical side by a rigid collagen-rich cuticle that serves as an exoskeleton. As in plant cells, this is likely to give the structural support that is provided by a highly organised cortical actin network in most animal cells (Salbreux et al., 2012). This may explain why cortical actin is not organized in the adult lateral epidermis. During moulting, however, when the cuticle is remodelled, a highly organized actin cytoskeleton comprised of circumferential bundles forms transiently, disappearing when the new cuticle is formed. This parallels the situation in the epidermis upon injury. Actin is rapidly assembled around the wound, and disassembled once it is repaired. Thus changes in membrane tension, either linked to alterations of the cuticle during development, or caused by a breach in its integrity are closely linked to cytoskeleton dynamics.

Wound repair has been previously suggested to involve a recapitulation of developmental programs (Martin and Parkhurst, 2004; Razzell et al., 2014). Thus for example, in the syncytial *Drosophila* early embryo, wound closure occurs through a myosin-II dependent purse-string mechanism, driven mainly by the contractile force exerted by myosin (Bement et al., 1999,Abreu-Blanco, 2011 #4455). This parallels the tight control exerted by myosin on the actin cytoskeleton during morphogenetic cell shape changes in *Drosophila* embryogenesis (Lecuit and Lenne, 2007). In *Xenopus* oocytes, if myosin motor activity is inhibited, however, wounds still close, even though at a much slower rate. The residual force driving wound closure is thought to be given by the treadmilling of actin, with a preferential recruitment of Rho GTPase towards the wound (Burkel et al., 2012).

Actin-dependent processes, including cell migration and cell protrusion, can be influenced by MT dynamics. The mechanisms that underlie their functional crosstalk are still being characterised (Dogterom and Koenderink, 2018). For example, a recent *in vitro* study showed that the MT + end-associated protein CLIP-170, together with the formin mDia1, can accelerate the actin filament polymerization which occurs at the tip of growing MTs (Henty-Ridilla et al., 2016). We showed here that in *C. elegans* the recruitment of MT + end binding protein EB1 to the wound precedes that of actin. We further demonstrated that perturbation of MT dynamics *in vivo* slows actin ring closure in *C. elegans* epidermis. Together, this suggests that MTs could facilitate actin ring closure. This MT-dependent mechanism, which is independent of myosin-based contractility (Xu and Chisholm, 2011), could arise because the epidermis is bounded by a rigid apical extracellular matrix, the tensile cuticle exoskeleton.

Injuring the epidermis not only leads to wound healing, but also provokes an innate immune response. Previous results in *C. elegans* suggested that the pathways involved in these 2 processes were distinct. Thus, although mutants for the p38 MAPK PMK-1 and for the SARM orthologue TIR-1, which acts upstream of the p38 cascade, are defective for the induction of AMP gene expression after wounding (Pujol et al., 2008a), they exhibit normal actin ring closure (Xu and Chisholm, 2011). If innate immune signal transduction pathway components are not required for wound repair, our results suggest, on the other hand, that cytoskeleton components can play a role in regulating AMP production. Indeed, several tubulin genes are required for AMP expression upon infection and wounding, potentially through their effect on the localization and dynamics of innate immune signalling proteins including SNF-12. The fact that overexpression in the epidermis of the MT severing protein Spastin caused a similar effect on SNF-12 dynamics as knocking down tubulin expression, supports a role for MT in positioning SNF-12-containing vesicles. Determining the precise nature of these vesicles, their connection with MTs and the mode of SNF-12 activation remain challenges for the future.

A further link between cytoskeleton reorganization and AMP production comes from the study of the protein DAPK-1, a negative regulator of both wound closure and innate immune gene expression (Tong et al., 2009). Interestingly, *dapk-1* mutants have defects in the epidermis that were shown to result from the hyperstabilization of MTs. DAPK-1 inhibits the function of PTRN-1, a MT - end binding protein, which acts as a MT nucleator and promotes MT stabilization. Loss of *ptrn-1* function suppresses the epidermal defects seen in *dapk-1* mutants as well as their constitutive AMP gene expression and accelerated wound closure. Loss of *dapk-1* function is associated with an increase in EB1 comets. This too is suppressed in a *dapk-1;ptrn-1* double mutant (Chuang et al., 2016). Taken together with our results, this suggests that the accelerated wound healing seen in *dapk-1* mutant worms may be due to an increased recruitment of EB1 protein thus speeding up MT-dependent actin ring closure.

The cytoskeleton is an assembly of co-regulating components that function together in a precise and dynamic way. Our results suggest that this co-regulation extends to the coordination of physical changes required for membrane wound healing and immune signal transduction driving transcriptional responses to injury.

## Material and Methods

### Nematode strains

All *C. elegans* strains were maintained on nematode growth medium (NGM) and fed with *E. coli* OP50, as described (Stiernagle, 2006): the wild type N2, IG274 *frIs7[col-19p::dsRed, nlp-29p::GFP]* IV (Pujol et al., 2008a), IG823 *frIs43[col-12p::SNF-12::GFP, ttx-3p::DsRed2]* and IG1235 *cdIs73[RME-8::mRFP, ttx-3p::GFP, unc-119(+)]; frIs43[col-12p::SNF-12::GFP, ttx-3p::DsRed2]* (Dierking et al., 2011), IG1327 *rde-1(ne219) V; juIs346[col-19p::RDE-1, ttx-3p::GFP]* III; *frIs7[nlp-29p::GFP, col-12p::dsRed]* IV (Zugasti et al., 2014), CZ13896 *juIs319[col-19p::GCaMP3, col-19p::tdTomato]* (Xu and Chisholm, 2011), CZ14453 *juEx3762[col-19p::EBP-2::GFP, ttx-3p::RFP]*, JLF302 *ebp-2(wow47[EBP-2::GFP]) II; zif-1(gk117) III* (Sallee et al., 2018), CZ14748 *juls352[GFP::moesin, ttx-3p::RFP]* I, CZ21789 *juSi239[col-19p::GFP::TBB-2]* I, CZ9334 *juEx1919[dpy-7p::GFP::RAB-5, ttx-3p::RFP]* (Chuang et al., 2016), SA854 *tbb-2(tj26[GFP::TBB-2])* III (Honda et al., 2017), SV1009 *Is[wrt-2p::GFP::PH-PLC1δ, wrt-2p::GFP::H2B, lin-48p::mCherry]* (Wildwater et al., 2011), MBA365 *Ex[dpy-7p::GFP::CAAX, myo-2p::GFP]* kindly provided by M. Barkoulas (UCL), RT343 *pwIs82[snx-1p::mRFP::SNX-1, unc-119(+)]*, RT424 *pwIs126[eea-1p::GFP::EEA-1, unc-119(+)]* (Shi et al., 2009), XW10992 *qxIs513[ced-1p::mCherry::PLC-1-PH]* and XW9653 *qxIs68[ced-1p::mCherry::RAB-7]* (Liu et al., 2012), ML1896 *mcIs35[lin-26p::GFP::TBA-2, pat-4p::CFP, rol-6(su1006)]; mcIs54[dpy-7p::SPAS-1_IRES_NLSmCherry, unc-119(+)]* X (Wang et al., 2015). All the transgenic strains carrying *frSi9* and *frSi13* (see below) were generated in this study (see Table S2 for a list of all strains).

### Constructs and transgenic lines

All the transgenic strains carrying *frIs7, frIs43, frSi9* or *frSi13* were obtained by conventional crosses and all genotypes were confirmed by PCR or sequencing. All constructs were made using Gibson Assembly (NEB Inc., MA).

*frSi9* is a single copy insertion on chromosome II at the location of the Nemagenetag Mos1 insertion (Vallin et al., 2012) *ttTi5605* of pNP151 (*col-62p::Lifeact::mKate_c-nmy3’utr*). pNP151 was obtained by insertion of the *col-62* promoter fused to Lifeact::mKate (Reymann et al., 2016) into the MosSCI vector pCFJ151 (Frokjaer-Jensen et al., 2008). It was injected into the EG6699 strain at 20 ng/μl together with pCFJ90 (*myo-2p::mCherry*) at 1.25 ng/μl, pCFJ104 (*myo-3p::mCherry*) at 5 ng/μl, pMA122 (*hsp16.41p::PEEL-1*) at 10 ng/μl, pCFJ601 (*eft-3p::Mos1 transposase*) at 20 ng/μl and pNP21 (*unc-53pB::GFP*) at 40 ng/μl. A strain containing the insertion was obtained following standard selection and PCR confirmation (Frokjaer-Jensen et al., 2008).

*frSi13* is a single copy insertion on chromosome II (*ttTi5605* location) of pNP159 (*dpy-7p::GFP::RAB-11*) by CRISPR using a self-excising cassette (SEC) (Dickinson et al., 2015). pNP159 was obtained by insertion of *dpy-7p::GFP::RAB-11* (kindly provided by Grégoire Michaux) into the pNP154 vector. pNP154 was made from a vector containing the SEC for single insertion on Chromosome II at the position of *ttTi5605* (pAP087, kindly provided by Ari Pani). pNP159 was injected in N2 at 10 ng/μl together with pDD122 (*eft-3p::Cas9*) at 40 ng/μl, pCFJ90 (*myo-2p::mCherry*) at 2.5 ng/μl, pCFJ104 (*myo-3p::mCherry*) at 5 ng/μl, and #46168 (*eef-1A.1p::CAS9-SV40_NLS::3’tbb-2* (Friedland et al., 2013)) at 30 ng/μl. Non-fluorescent roller worms were selected then heat shocked to remove the SEC by FloxP as described in (Dickinson et al., 2015). Plasmid and primers sequences are available upon request.

Different transgenic strains containing several tagged version of SNF-12 were generated including IG1784 *frEx597[pSO16(col-12p::SNF-12::mKATE_3’unc-54), pCFJ90(myo-2p::mCherry]*, IG1270 *frEx453[pMS8(col-12p::GFP::STA-2), pMS9(col-12p::mCherry::SNF-12)]* and IG1663 *frEx577[pNP158(snf-12p::SNF-12::GFP_3’snf-12, pCFJ90(myo-2p::mCherry), pCFJ104(myo-3p::mCherry)];* all strains are given in Table S2.

### Infection and needle wound

Eggs prepared by the standard bleach method were allowed to hatch in 50 mM NaCl in the absence of food at 25 °C. Synchronized L1 larvae were transferred to NGM agar plates spread with *E. coli* OP50 and cultured at 25 °C until the L4 stage (40 hours) before being exposed to fungal spores on standard NGM plate spread with OP50, as previously described (Pujol et al., 2001). To count the number of spores, young adult worms were infected for 8 hours at 25 °C then observed on a slide under the microscope. Statistical significance was determined using ANOVA Bonferoni’s test (Graphpad Prism). Needle wounding was performed as previously described (Pujol et al., 2008a) with a standard microinjection needle under a dissecting microscope by pricking the worm posterior body or tail on agar plates; worms were analysed after 6h.

### Fluorescent reporter analyses

Analysis of *nlp-29*p::GFP expression was quantified with the COPAS Biosort (Union Biometrica; Holliston, MA) as described in (Pujol et al., 2008a). In each case, the results are representative of at least 3 independent experiments with more than 70 worms analysed. The fluorescent ratio between GFP and size (time of flight; TOF) is represented in arbitrary units. Statistical significance was determined using a non-parametric analysis of variance with a Dunn’s test (Graphpad Prism). Fluorescent images were taken of transgenic worms mounted on a 2 % agarose pad on a glass slide anesthetized with 0.01 % levamisole, using the Zeiss AxioCam HR digital colour camera and AxioVision Rel. 4.6 software (Carl Zeiss AG).

### RNAi

The RNAi bacterial clones were obtained from the Ahringer library (Kamath et al., 2003) or Vidal library (Rual et al., 2004) and sequenced to confirm their identity using CloneMapper (Thakur et al., 2014). RNAi feeding experiments were performed at 25 °C and the RNAi clone *sta-1* was used as a negative control. To avoid developmental delay or lethality, RNAi experiments of tubulin or actin genes were either performed from L4 stage or in IG1327 (*rde-1(ne219) V; juIs346[col-19p::RDE-1, ttx-3p::GFP]* III; *frIs7[nlp-29p::GFP, col-12p::dsRed]* IV) worms (Zugasti et al., 2016) allowing gene silencing specifically in the epidermis from the L4 stage.

### Image acquisition and laser wound

Since in preliminary experiments we observed that immobilization using latex beads impacted vesicle and protein dynamics in the epidermis, young adult worms (containing less than 5 eggs) were immobilized on a slide using 0.01 % levamisole. Time lapse and/or Z stack were acquired using two different spinning disk microscopes: an inverted Visitron Systems GmbH spinning disc with a Nikon 40X oil, 1.3 NA objective and 1.5X lens, and a Roper inverted spinning disc with a Nikon 100X oil, 1.4 NA objective. The two systems acquire images using Visitron and MetaMorph, respectively. Laser wounding was performed by using an imaging 405 nm laser (22 mW) operating in a FRAP mode, using either Visitron or iLas FRAP Systems, respectively. 1-3 sec of laser pulses were applied on a circular region of interest (ROI) of 21 × 21 pixels with the Nikon (~1.68 μm in diameter) or 8 × 8 pixels with the Visitron (~2.13 μm in diameter) on the apical plane of the lateral epidermis either left or right of the seam cells. Each worm was wounded from 1 to 3 times, if possible sequentially on opposite sides of the seam cells. Time lapse acquisitions were 15 to 20 min long. Each Z stack was acquired every 15 sec and the Z size was 0.3 or 0.5 μm. High temporal resolution was obtained on one single plane with a time interval of 0.2 to 0.5 ms. Laser wound was performed between the first and the second Z stack. For colocalization analysis, we initially controlled that there was no signal cross talk using single labelled strains. We then performed sequential acquisitions on the strains expressing 2 fluorescent proteins. Experiments requiring comparative analysis were performed on the same day.

### Image analysis

All image processing was done using Fiji software. A custom Image J macro was used to correct for focus z-drift during the 20 min acquisitions. XY drift was corrected either with the “Image Stabilizer” plugin from ImageJ or the online available macro “NMSchneider/FixTranslation”.

To analyse EB1 density before wounding, we manually counted fluorescent dots as described in (Chuang et al., 2016), in a 300 μm^2^ ROI in the lateral hyp7 or in an 80 μm^2^ ROI in the seam cell in 5 worms per RNAi condition. Similarly, we measured SNF-12 cluster density in the lateral hyp7, analysing 6 worms per RNAi condition. Quantification of EB1 recruitment at the wound site was performed by measuring the RawIntDen (sum of the pixel intensities) of a selected ROI of 256 μm^2^ for EB1 centred at the wound site. Since EB1 brightness differed before wounding between worms, for each single wound, only pixels with values above a defined brightness threshold were considered, the threshold being set to the [mean + 3 × standard deviation] in the ROI before wounding. Quantification of SNF-12 recruitment was performed by measuring the mean intensity of a selected ROI of 18 μm^2^ centred at the wound site at specific time points. Statistical significance was determined using a non-parametric Mann-Whitney test (Graphpad Prism).

To visualise the speed and directionality of EB1, RAB-11 and SNF-12, a temporal projection was done using a temporal-Color code (plugin from Kota Miura) on 120 frames over 2 min (1 frame per second) before and on 32 frames over 10 min after wounding for SNF-12. To analyse the speed of recruitment of EB1 versus actin, the radial profile (plugin from Paul Baggethun) was calculated, after determination of the centroid of the wound for each time point by manually selecting the wound area. For each radial profile, we extracted the radius corresponding to the maximum intensity and plotted this radius versus time.

To analyse particle motility before and after wounding, we used the macro KymographClear described in (Mangeol et al., 2016). For EB1, RAB-11 and SNF-12, we acquired movies with time interval of 300 ms, 400 ms and 200 ms, respectively. Since SNF-12 was observed to move much slower than the other dynamic proteins studied, a substack with a time interval of 20 s was generated. SNF-12 quantifications were done for particles with a minimum displacement of 5 pixels within 120 sec for before wounding and within either 210 or 420 sec for after wounding.

To analyse actin ring closure, an ROI of 80 pixels^2^ located on the centre of the wound was selected 15 mn post wounding. The area inside the actin ring was detected using the Huang automatic threshold method from Image J.

## Supporting information

Plasma membrane rapidly reorganizes at the wound site. Imaging Ex dpy-7p::GFP::CAAX; * indicates the wound site; [min:sec].

Rapid PIP2 domain reorganization at the wound site. Imaging Is wrt-2p::GFP::PH-PLC1??; * indicates the wound site; [min:sec].

Actin reorganization at the wound site. Actin ring closure within 20 min. Imaging Si col-62p::Lifeact::mKate strain; * indicates the wound site; [min:

EB1/EBP-2, a plus end MT binding protein, is recruited at the wound site.

Cytoskeleton reorganization at the wound site. Actin (in red) and MT plus ends (in green) form 2 concentric rings with MT forming the inner array.

Microtubules regrow around the wound site. Imaging: KI tbb-2p::TBB-2::GFP strain; * indicates the wound site; [min:sec].

Microtubules regrow around the wound site in front of actin. Imaging: KI tbb-2p::TBB-2::GFP strain; Si col-62p::Lifeact::mKate; * indicates the wound

SNF-12 dynamics in adult epidermis. Imaging Is col-12p::SNF-12::GFP strain; [min:sec].

SNF-12 vesicles are recruited at the wound site. This movie represents SNF-12 recruitment from 6 min and 54 sec post wound

RAB-5 dynamics in adult epidermis. Imaging juEx1919 dpy-7p::GFP::RAB-5 strain.* indicates the wound site; [min:sec].

RAB-5 recruitment at the wound site. Imaging juEx1919 dpy-7p::GFP::RAB-5 strain; * indicates the wound site; [min:sec].

RAB-11 dynamics in adult epidermis. Imaging ; frSi13 dpy-7p::GFP::RAB-11 strain; * indicates the wound site; [min:sec].

RAB-11 recruitment at the wound site. Imaging frSi13 dpy-7p::GFP::RAB-11 strain; * indicates the wound site; [min:sec].

## Acknowledgement

We thank Annie Bonnet, Vincent Rouger, Guillaume Bordet and Pierre Mangeol for their contributions, Asako Sugimoto, Andrew Chisholm, Michel Labouesse, Sander van den Heuvel, Michalis Barkoulas, Bart Grant, Xiaochen Wang, Jessica Feldman and the *Caenorhabditis* Genetics Center (University of Minnesota, Minneapolis, MN) supported by the National Institutes of Health Office of Research Infrastructure Programs (P40 OD010440) for strains, Ari Pani, Anne Cecile Rayman, Gregoire Michaux and Michael Sixt for plasmids, Chris Crocker at Wormatlas (supported by NIH OD010943 to David Hall) for diagrams, and Didier Marguet, Ariane Abrieu, Alphée Michelot, Thomas Lecuit and members of our lab for discussions and critical reading of the manuscript. Worm sorting was performed by Jerome Belougne using the facilities of the French National Functional Genomics platform, supported by the GIS IBiSA and Labex INFORM. We thank the imaging core facility (ImagImm) of the Centre d’Immunologie de Marseille-Luminy (CIML) supported by the French National Research Agency program (France-BioImaging).

## Fundings

This work was supported by the French National Research Agency: ANR-16-CE15-0001-01, ANR-12-BSV3-0001-01, ANR-11-LABX-0054, ANR-11-IDEX-0001-02, ANR-10-INBS-04-01 (France Bio Imaging), CNRS, Aix Marseille University, National institute of Health and Medical Research (Inserm).

---

**Table S1.** MT-related genes identified in a genome-wide screen for regulators of AMP gene expression from. (Zugasti et al., 2016).

| **WormBase Gene ID**<br>**(WS266)** | **Public Name** | **Sequence Name** | **Description** |
| --- | --- | --- | --- |
| WBGene00000962 | *dhc-1* | T21E12.4 | dynein heavy chain homolog |
| WBGene00001017 | *dnc-1* | ZK593.5 | dynactin complex subunit p150 homolog |
| WBGene00001832 | *hcp-4* | T03F1.9 | centromere protein (CENP)-C homolog |
| WBGene00002231 | *knl-1* | C02F5.1 | essential kinetochore component |
| WBGene00002845 | *let-711* | F57B9.2 | NOT1 orthologue |
| WBGene00004274 | *rab-11.1* | F53G12.1 | small GTPase homologous to the Rab GTPases |
| WBGene00010556 | *rack-1* | K04D7.1 | orthologue of vertebrate Receptor for Activated C Kinase |
| WBGene00006529 | *tba-2* | C47B2.3 | alpha-tubulin. |
| WBGene00006530 | *tba-4* | F44F4.11 | alpha tubulin |

**Table S2.**
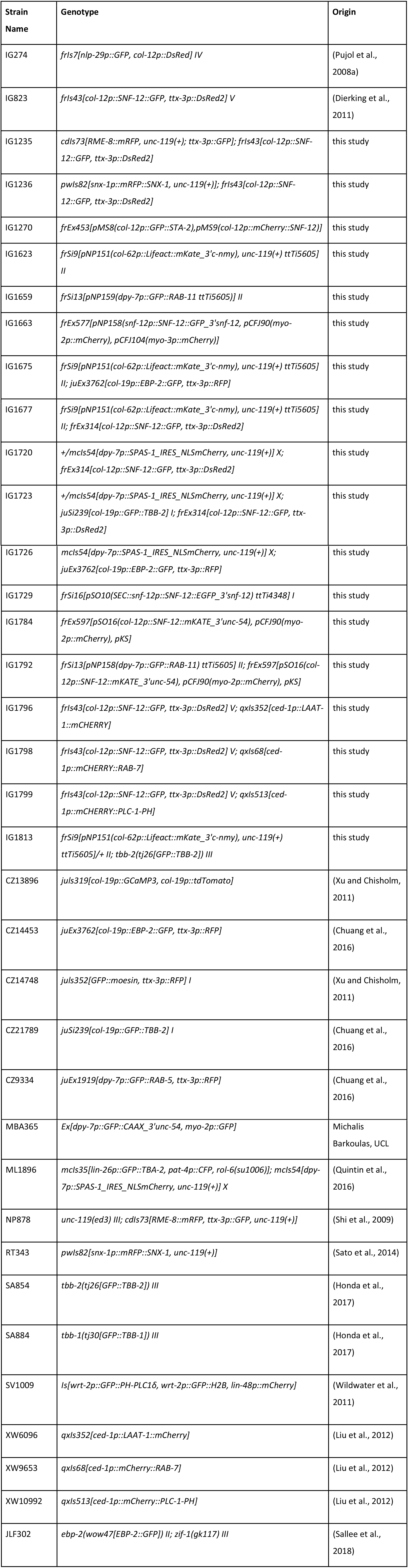
List of the *C. elegans* strains used in this work.

**Fig. S1:**
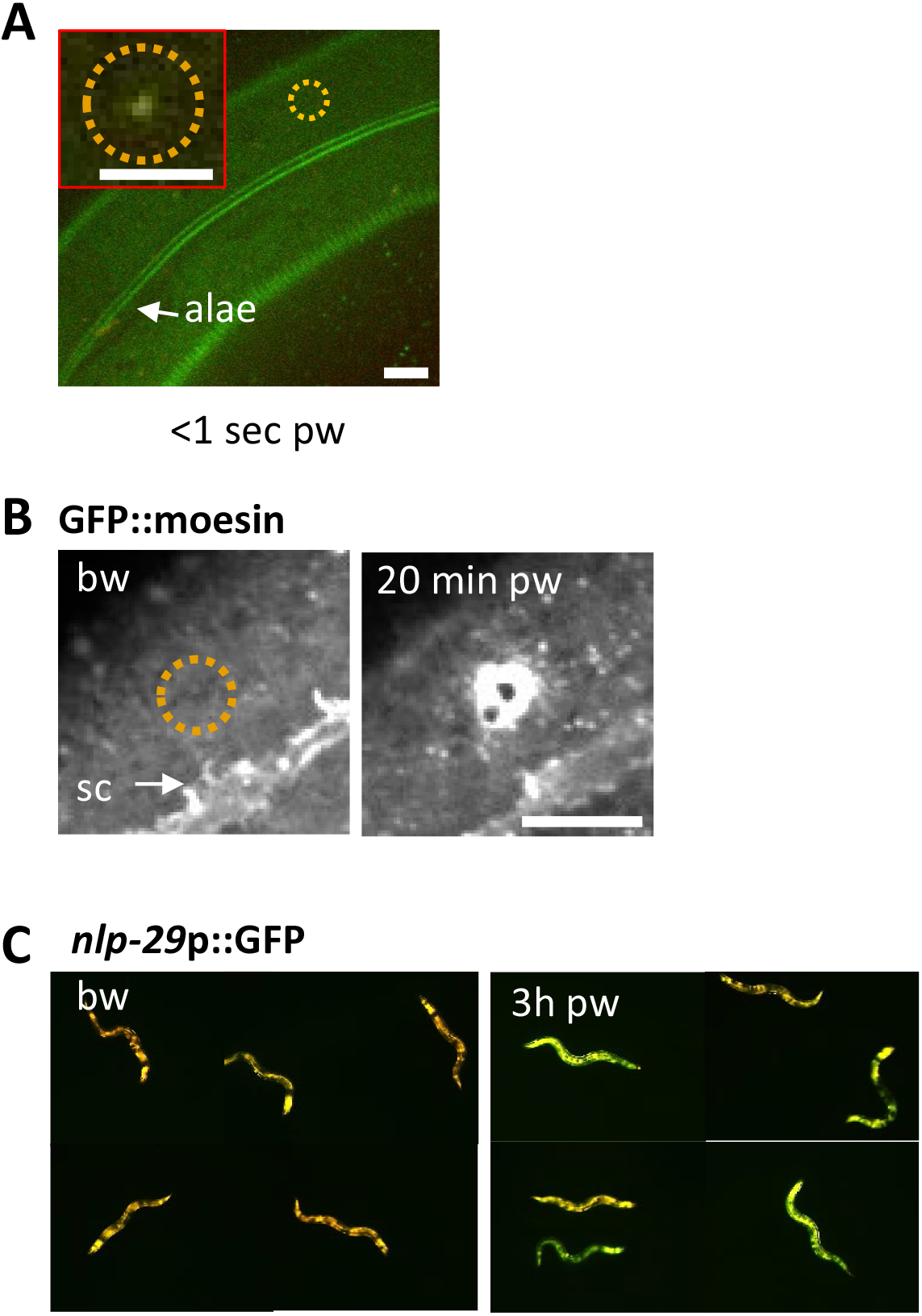
Reproduction of known wound hallmarks following injury using a 405 nm laser. Wounding *C. elegans* lateral epidermis with a 405 nm laser causes immediate formation of an autofluorescent scar (A) and later an actin ring, revealed with the actin-binding protein GFP::moesin (B) at the wound site. Representative images of wild-type (A) and *col-19p::GFP::moesin* (B) worms. The dashed circle is centred on the wound site; bw, before wound; pw, post wound; sc, seam cells. Scale bar 10 μm except in the inset 5 μm. (C) 3 hours after wounding, young adult worms exhibit an increased expression of a *nlp-29p::GFP* reporter, a read out of the immune response (right), compared to unwounded worms (left). The worms express dsRed constitutively in the epidermis. Red and green fluorescence are visualized here simultaneously.

**Fig. S2:**
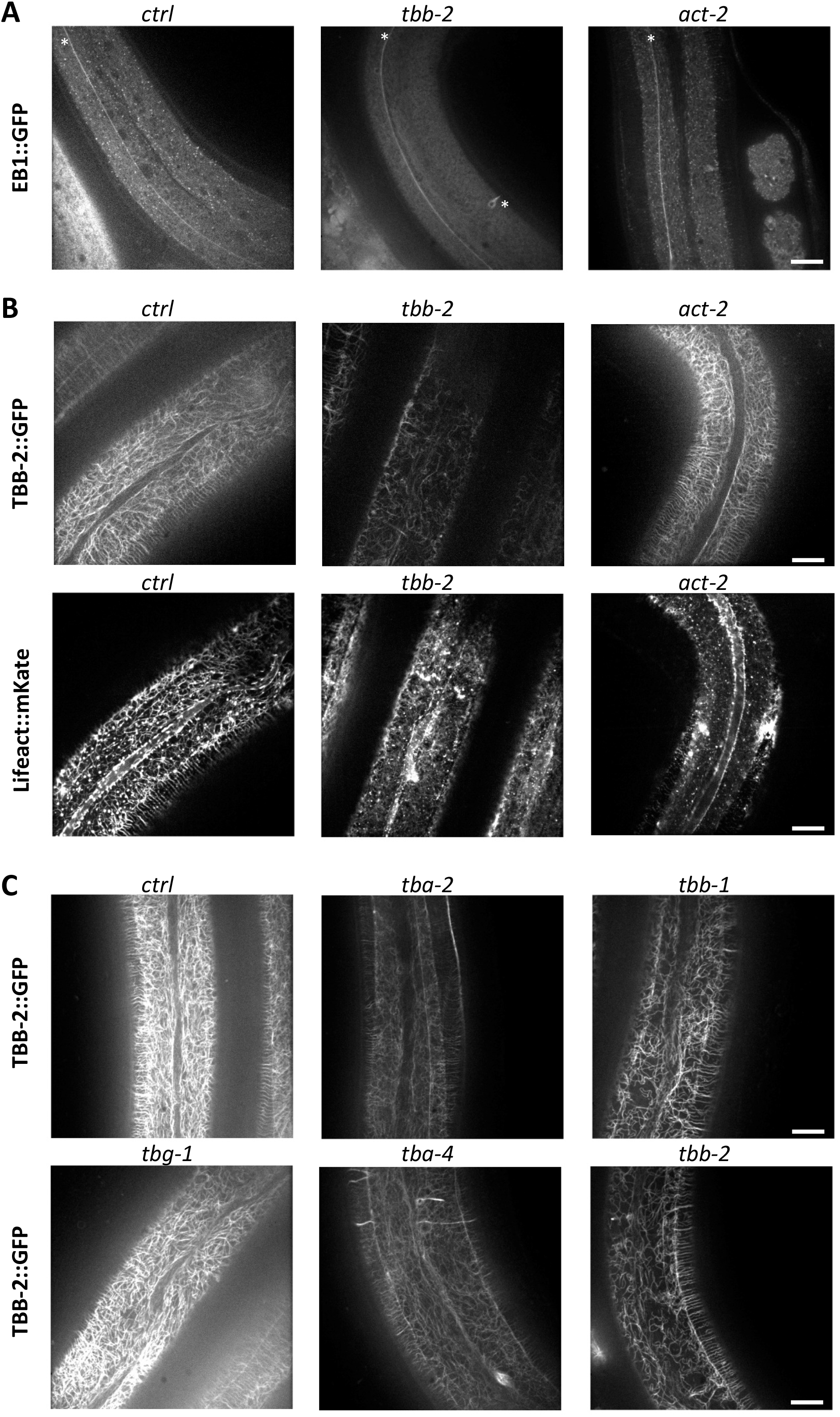
Tubulin is required for non-centrosomal MT organisation and dynamics in the lateral epidermis. TBB-2 but not ACT-2 is required for the normal pattern of EB1 comets (A) and MTs (B). Representative images of (A) *EBP-2::GFP KI* worms (white star point to mechanosensory axons), or (B) TBB-2::GFP (top) and Lifeact::mKate (bottom) transgenes, after short temporal inactivation of control (*sta-1; ctrl), tbb-2* and *act-5* genes; scale bar 10 μm. (C) RNAi against *tba-2, tba-4, tbb-1* and *tbb-2*, but not *tbg-1* alters the pattern of MTs in the adult epidermis. Representative images of worms expressing TBB-2::GFP after short temporal inactivation of a *ctrl, tba-2, tba-4, tbb-1, tbb-2* and *tbg-1* genes; scale bar 10 μm.

**Fig. S3:**
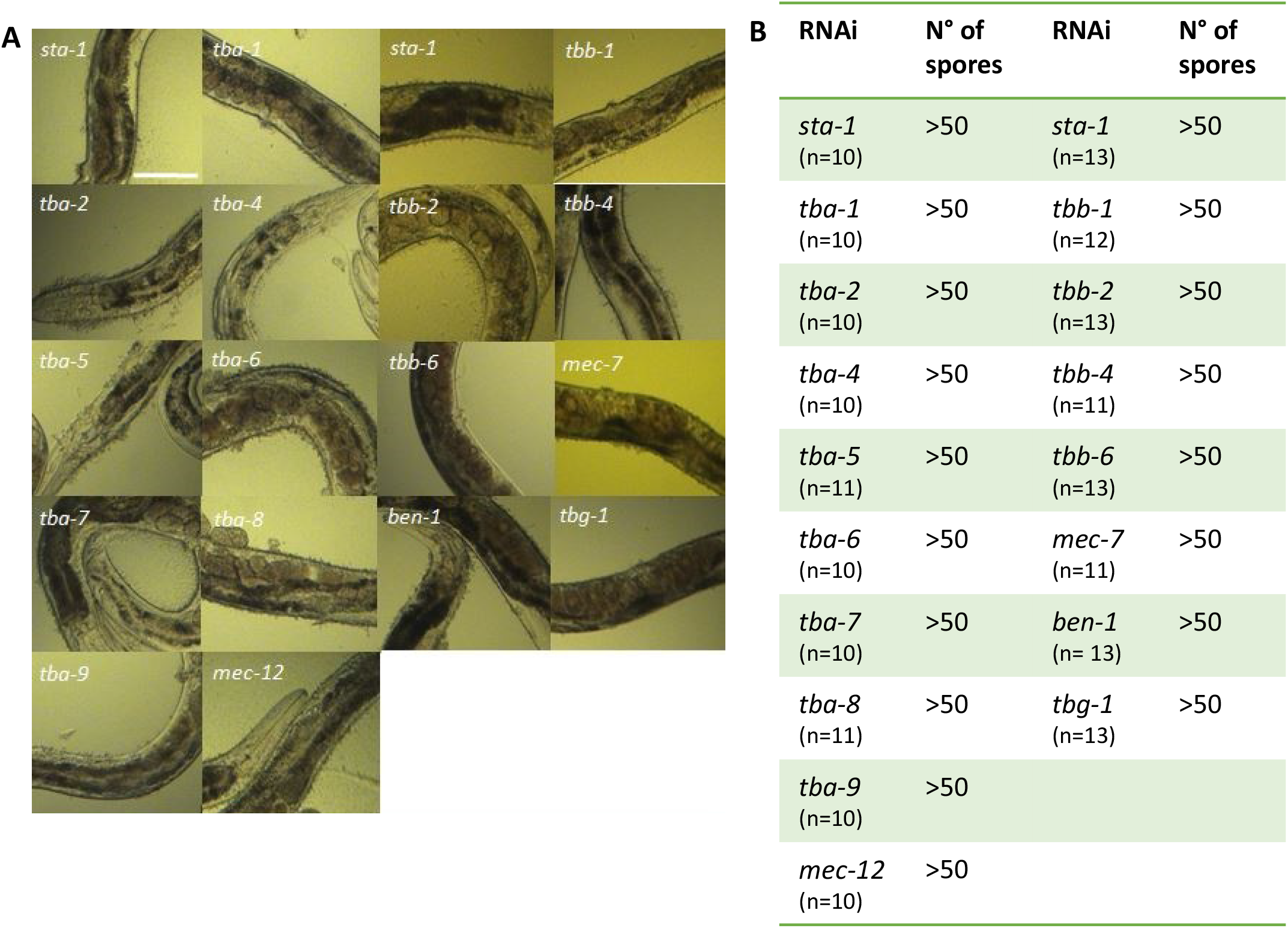
Worms are efficiently infected after knock-down of tubulin gene expression by RNAi. (A) Representative white-field images of IG1327 worms after infection with *D. coniospora* for 18 h. Before infection worms were fed with RNAi clones against different tubulin-α, β and γ isoforms. Each RNAi clone was tested 4 times. Scale bar 100 μm. (B) Table summarising the mean value of the number of spores adhering to the infected worms, after 18 h of *D. coniospora* infection. IG1327 *rde-1(ne219) V; juIs346[col-19p::RDE-1, ttx-3p::GFP] III; frIs7[nlp-29p::GFP, col-12p::dsRed] IV*

**Fig. S4:**
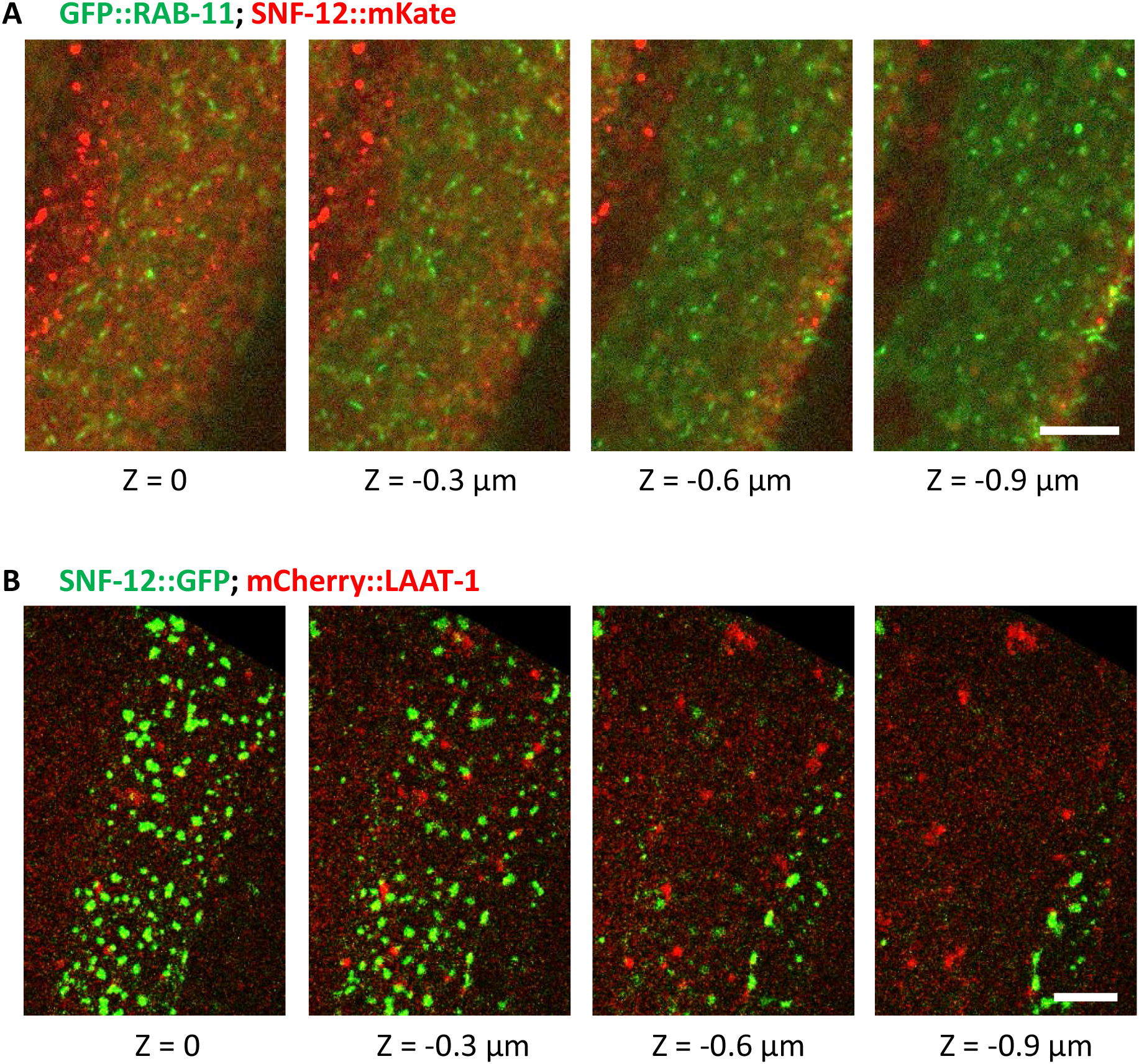
SNF-12 does not colocalize with RAB-11 or LAAT-1. Colocalization analyses to compare SNF-12 localization with known vesicular markers. Recycling endosomes (A) and lysosomes (B) are visualized as RAB-11 (A) and LAAT-1 (B) positive vesicles, respectively. SNF-12 did not colocalize with these markers. SNF-12 was in the same focal plane as the most apical RAB-11 recycling endosomes (A) and apical to LAAT-1 positive lysosomes (B). Z indicates the different focal planes. Representative images of *dpy-7p::GFP::RAB-11; col-12p::SNF-12::mKate* (A) and *col-12p::SNF-12::GFP; ced-1p::LAAT-1::mCherry* (B). Scale bar 5 μm.

**Fig. S5:**
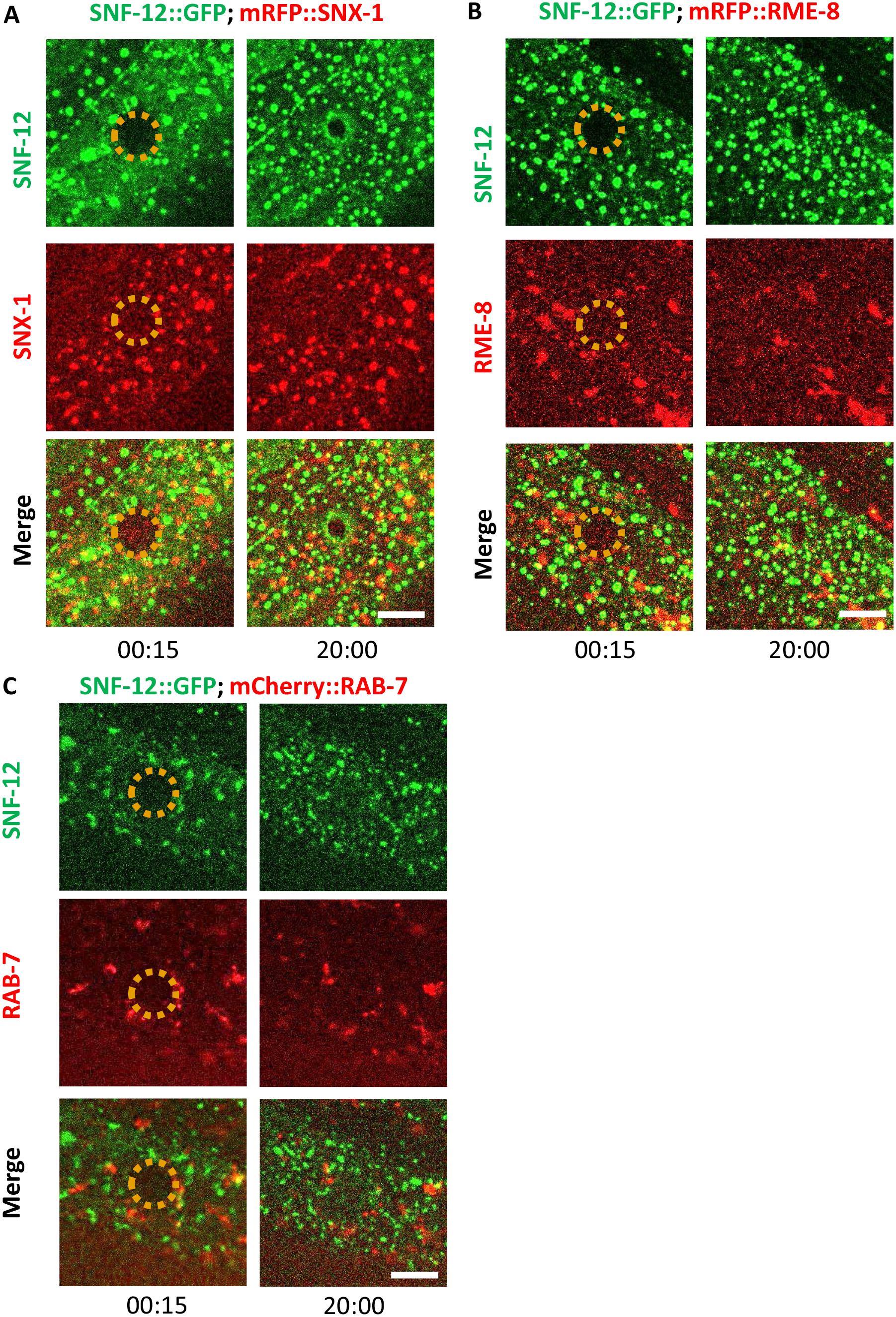
SNF-12 exhibits a specific dynamic behaviour upon wounding. Unlike early endosomes (A-B) or late endosomes (C) visualized with SNX-1 (A), RME-8 (B) and RAB-7 (C) reporter proteins, respectively, upon wounding, SNF-12 clusters are locally recruited to the wound site. Representative figures of *col-12p::SNF-12::GFP; snx-1p::mRFP::SNX-1* (A), *col-12p::SNF-12::GFP; RME-8::mRFP* (B) and *col-12p::SNF-12::GFP; ced-1p::mCherry::RAB-7* worms (C). The dashed circle is centred on the wound site; time post wound [min:sec]; scale bar 5 μm.

**Fig. S6:**
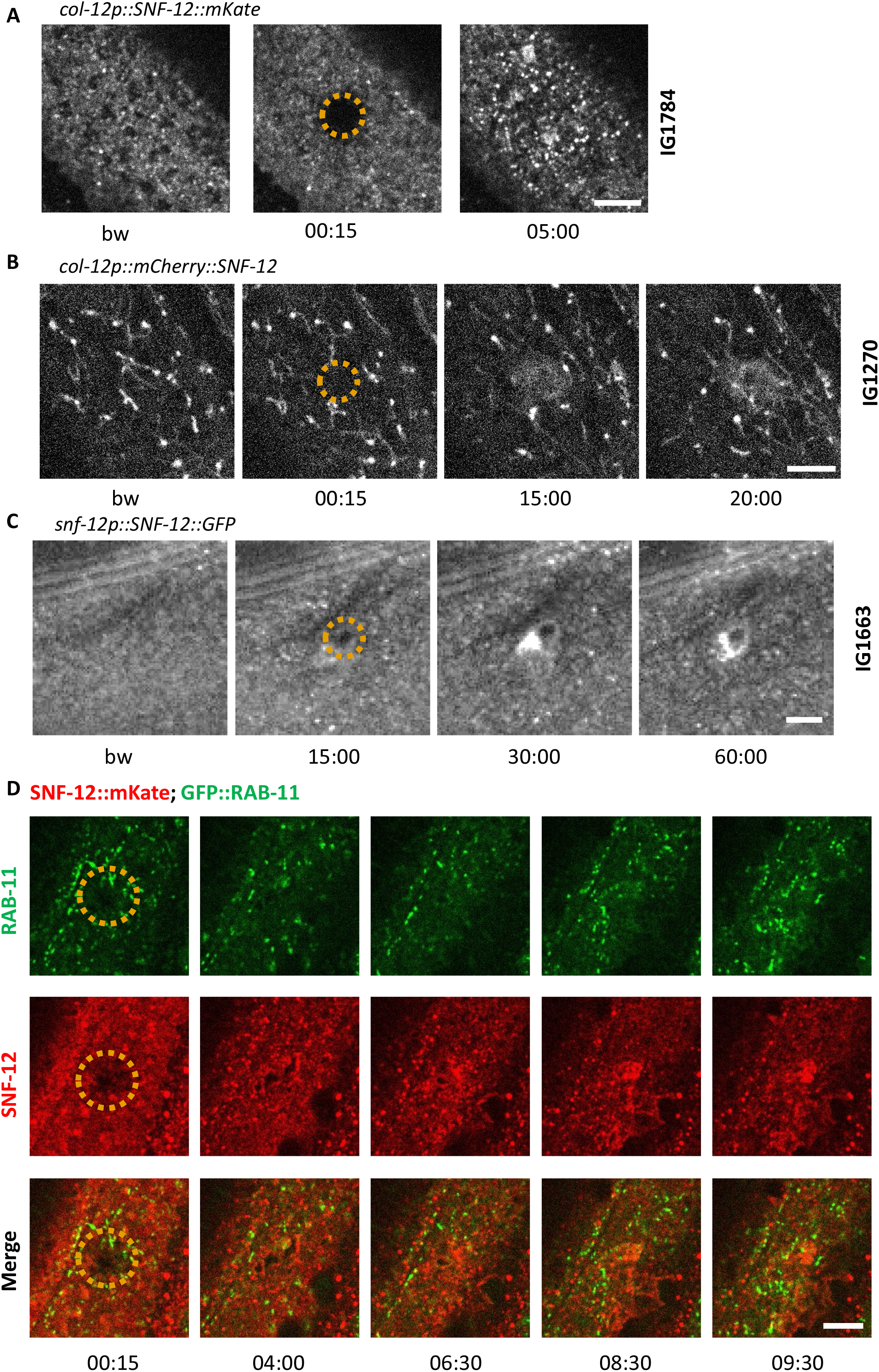
SNF-12 recruitment towards the wound site is observed using different chimeric reporter proteins. Representative images of worms carrying *col-12p::SNF-12::mKate* (IG1784; A), *col-12p::mCherry::SNF-12* (IG1270; B), *snf-12p::SNF-12::GFP* (IG1663; C) transgenes. In (B), the large aggregates and filaments that are observed in addition to the SNF-12-positive clusters are probably artefacts due to the known propensity of mCherry to aggregate. The dashed circle is centred on the wound site; time post wound [min:sec]; scale bar 5 μm. (D) Recruitment of SNF-12 and RAB-11 at the wound results in clusters that do not greatly overlap.

**Fig. S7:**
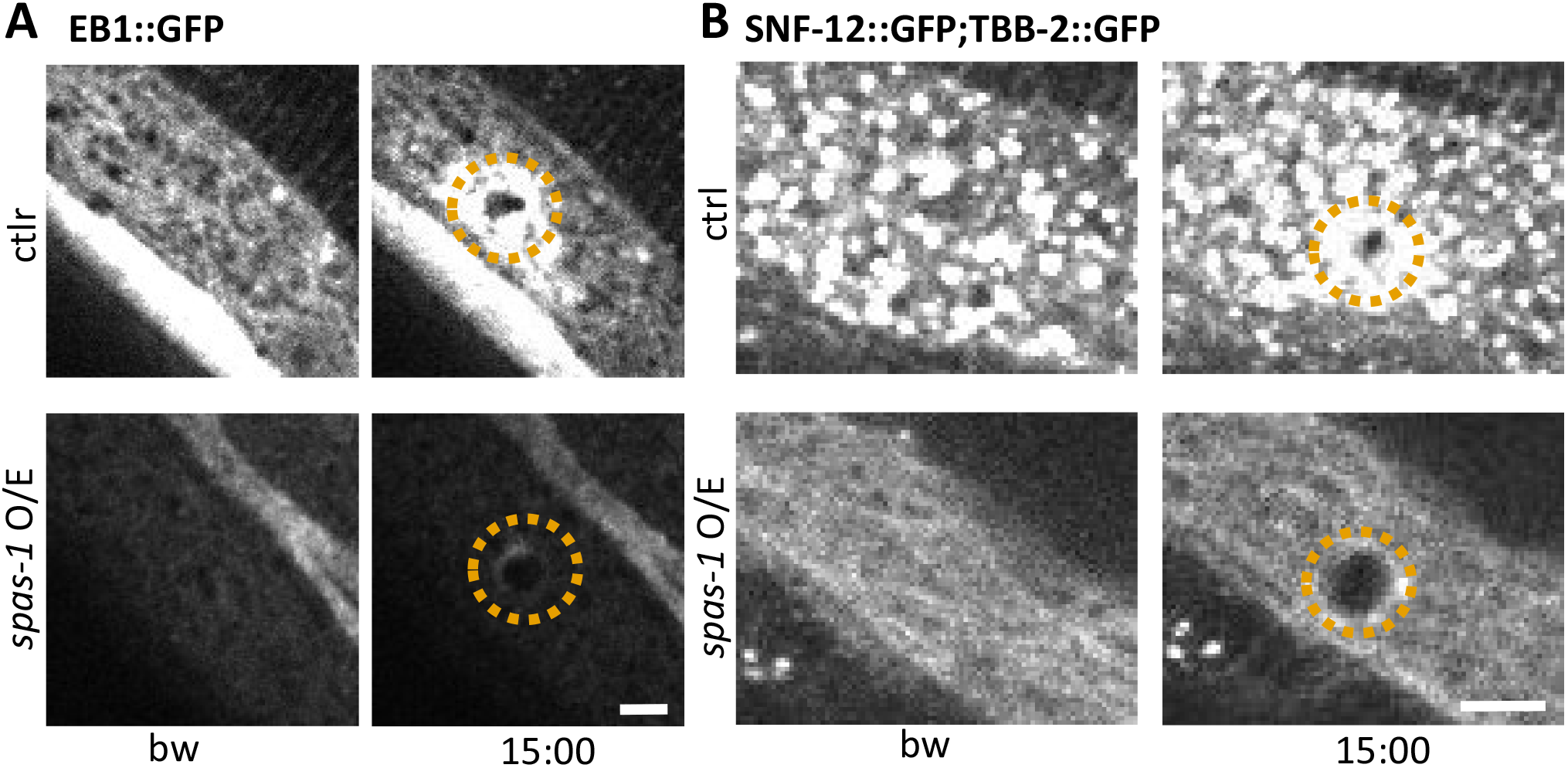
Spastin-1 overexpression drastically alters MT and SNF-12 dynamics before and after wounding. Overexpression of the severing protein Spastin (SPAS-1) in the epidermis under the control of the *dpy-7* promoter (*spas-1 O/E*) affects both EB1 and SNF-12 patterning and recruitment towards wounds. Representative images of worms carrying *col-19p::EBP-2::GFP* (A) and *col-12p::SNF-12::GFP; col-19p::GFP::TBB-2* (B) without and with the *dpy-7p::SPAS-1* construct (upper and lower panels, respectively) before and after wounding. The dashed circle is centred on the wound site; time post wound [min:sec]; scale bar, 5 μm.

## Movies

**Movie 1: Plasma membrane rapidly reorganizes at the wound site**.

Imaging Ex *dpy-7p::GFP::CAAX*; * indicates the wound site; [min:sec].

**Movie 2: Rapid PIP_2_ domain reorganization at the wound site**.

Imaging Is *wrt-2p::GFP::PH-PLClδ*; * indicates the wound site; [min:sec].

**Movie 3: Actin reorganization at the wound site**.

Actin ring closure within 20 min. Imaging Si *col-62p::Lifeact::mKate* strain; * indicates the wound site; [min:sec].

**Movie 4: EB1/EBP-2, a plus end MT binding protein, is recruited at the wound site**.

This movie represents a different wound from the one in Fig. 3B. Imaging Ex *col-19p::EBP-2::GFP* strain; * indicates the wound site; [min:sec].

**Movie 5: Cytoskeleton reorganization at the wound site**.

Actin (in red) and MT plus ends (in green) form 2 concentric rings with MT forming the inner array.

This movie represents a different wound from the one in Fig. 3D. Imaging Si *col-62p::Lifeact::mKate*; Ex *col-19p::EBP-2::GFP* strain; * indicates the wound site; [min:sec].

**Movie 6: Microtubules regrow around the wound site**.

Imaging: *KI tbb-2p::TBB-2::GFP* strain; * indicates the wound site; [min:sec].

**Movie 7: Microtubules regrow around the wound site in front of actin**.

Imaging: *KI tbb-2p::TBB-2::GFP* strain; Si *col-62p::Lifeact::mKate*; * indicates the wound site; [min:sec].

**Movie 8: SNF-12 dynamics in adult epidermis**.

Imaging Is *col-12p::SNF-12::GFP* strain; [min:sec].

**Movie 9: SNF-12 vesicles are recruited at the wound site**.

This movie represents SNF-12 recruitment from 6 min and 54 sec post wound and it is a different wound from the one shown in Fig. 6C.

Imaging Is *col-12p::SNF-12::GFP* strain; * indicates the wound site; [min:sec].

**Movie 10: RAB-5 dynamics in adult epidermis**.

Imaging *juEx1919 dpy-7p::GFP::RAB-5* strain.* indicates the wound site; [min:sec].

**Movie 11: RAB-5 recruitment at the wound site**.

Imaging *juEx1919 dpy-7p::GFP::RAB-5* strain; * indicates the wound site; [min:sec].

**Movie 12: RAB-11 dynamics in adult epidermis**.

Imaging *frSi13 dpy-7p::GFP::RAB-11* strain; * indicates the wound site; [min:sec].

**Movie 13: RAB-11 recruitment at the wound site**.

Imaging *frSi13 dpy-7p::GFP::RAB-11* strain; * indicates the wound site; [min:sec].

